# Metagenomic, metabolomic, and lipidomic shifts associated with fecal microbiota transplantation for recurrent *Clostridioides difficile* infection

**DOI:** 10.1101/2024.02.07.579219

**Authors:** Arthur S McMillan, Guozhi Zhang, Michael K Dougherty, Sarah K McGill, Ajay S Gulati, Erin S Baker, Casey M Theriot

## Abstract

Recurrent *C. difficile* infection (rCDI) is an urgent public health threat for which the last resort and lifesaving treatment is a fecal microbiota transplant (FMT). However, the exact mechanisms which mediate a successful FMT are not well understood. Here we use longitudinal stool samples collected from patients undergoing FMT to evaluate changes in the microbiome, metabolome, and lipidome after successful FMTs. We show changes in the abundance of many lipids, specifically acylcarnitines and bile acids, in response to FMT. These changes correlate with Enterobacteriaceae, which encode carnitine metabolism genes, and Lachnospiraceae, which encode bile salt hydrolases and *baiA* genes. LC-IMS-MS revealed a shift from microbial conjugation of primary bile acids pre-FMT to secondary bile acids post-FMT. Here we define the structural and functional changes in successful FMTs. This information will help guide targeted Live Biotherapeutic Product development for the treatment of rCDI and other intestinal diseases.

## Introduction

*Clostridioides difficile* is a Gram-positive, anaerobic, spore forming pathogen that is a major cause of morbidity and remains a leading healthcare-associated infection ^1–3^. One of the most significant risk factors for acquiring *C. difficile* infection (CDI) is antibiotic usage. Antibiotic usage alters the protective native gut microbiota leading to a loss of colonization resistance against *C. difficile* ^4–6^. Once colonization resistance decreases, *C. difficile* spores are able to germinate and vegetative cells can colonize the gut, ultimately producing toxin resulting in a range of clinical disease including diarrhea, pseudomembranous colitis and even death ^7,8^.

Current recommendations for the treatment of CDI are a course of antibiotics, either vancomycin or fidaxomicin, which continue to alter colonization resistance and can result in a re-emergence of CDI ^4,9^. This re-emergence after initial clinical cure of the baseline episode is called recurrent *C. difficile* infection (rCDI) ^9,10^. rCDI occurs in up to ∼35% of patients after cessation of their first course of antibiotics, and each additional antibiotic treatment results in an increased chance of future recurrence ^11,12^. Fecal microbiota transplantation (FMT) is a last resort guideline-recommended therapy for rCDI associated with cure rates around 80-98%, where the stool from healthy donors is introduced into patients with rCDI ^9,13–16^. FMTs are the last line of treatment for patients with recurrent rCDI, however they are not standardized and may have unknown long-term health consequences as seen by the deaths due to the transfer of donor stool with antibiotic resistant bacteria acquired from a stool bank ^17^. In 2023, FDA approved the first microbiota focused therapeutic or Live Biotherapeutic Products (LBP) (Rebyota and Vowst) to treat rCDI. These products attempt to standardize the donor feces to make the FMT safer, however these products are still derived from raw stool ^18,19^. While the microbiome field has progressed greatly, the mechanisms mediating a successful FMT are still unknown ^20,21^. Thus, determining molecularly and mechanistically how an FMT clears rCDI in patients is essential in order to design safer, targeted bacterial therapeutics to prevent and treat both primary and rCDI.

To date, multiple studies have reported structural changes to the microbiota in rCDI after FMT ^14,22–26^. Specifically, they have observed an increase in the number of commensal bacteria from the Phyla of Firmicutes and Bacteroidetes, and a decrease in Proteobacteria ^27^. These changes are always accompanied with an increase in alpha diversity ^14,22–26^. While it is not known why an increase in alpha diversity allows for a successful FMT, a recent study found an increase in diversity were able to exclude both *Klebsiella pneumoniae* and *Salmonella enterica* serovar Typhimurium from colonizing the gut through the mechanism of nutrient blocking, by consuming nutrients the pathogen needed ^28^. Members of the gut microbiota also produce and consume the same amino acids that *C. difficile* requires for growth. Recently, donor stool used in successful FMTs was found to contain high levels of amino acid biosynthesis genes suggesting amino acid cross-feeding as a potential mechanism ^29^. *C. difficile* requires amino acids for which it is auxotrophic, including cysteine, isoleucine, leucine, proline, tryptophan, and valine. Moreover, it prefers Stickland fermentation substrates, such as isoleucine, leucine, glycine, proline, and hydroxyproline ^30–32^. The newly introduced donor stool may have a microbial community able to compete for nutrients with *C. difficile* that is reflected as an increase in alpha diversity associated with FMT for rCDI.

Changes to the lipidome were also observed alongside the microbiota after FMT for rCDI. ^33–35^. FMT and LBP induced changes to the gut lipidome in rCDI patients, specifically an increase in secondary bile acids and short chain fatty acids (SCFAs), which are able to negatively impact *C. difficile* growth *in vitro* ^19,34–39^. *C. difficile* is exquisitely sensitive to bile acids ^37^, particularly secondary bile acids, which are made by gut microbial enzymes from host derived primary bile acids. Gut bacteria drive changes to the bile acid pool through a variety of mechanisms, including deconjugation of bile acids by the removal of the conjugated amino acid via bile salt hydrolases (BSHs), and the generation of secondary bile acids by the removal of the 7α-hydroxyl group via the *bile acid inducible* (*bai*) operon. These enzymes are associated with successful FMTs in both human and mouse models ^25,40–42^. BSHs are known to deconjugate either glycine or taurine from host conjugated bile acids. However, recent studies suggest that BSHs have the ability to reverse this reaction and reconjugate bile acids with a variety of other amino acids thereby creating microbially conjugated bile acids (MCBAs), also referred to as bacterial bile acid amidates (BBAAs) ^43,44^. MCBAs have previously been identified in a small sample set of rCDI patients treated with FMT by our group ^45^. In a recent study, a BSH cocktail given to mice increased MCBAs and was able to restrict *C. difficile* growth *in vivo* ^46^. Furthermore, certain MCBAs also inhibit germination, vegetative growth, and toxin expression of *C. difficile in vitro* ^46^. Thus, the return of a diverse community that restores secondary bile acid metabolism could be detrimental to *C. difficile* and also play a role in mediating FMT success. Further mechanistic studies paired with more sensitive metabolomic platforms are needed to determine what constitutes a successful FMT for the treatment of rCDI.

In this study, we utilize a multi-omic approach, leveraging metagenomics and multiple metabolomic analyses that range in sensitivity, to identify changes in the microbiome, metabolome and lipidome after a successful FMT in patients with rCDI. We observe significant changes in the microbiota structure and lipid metabolism between pre- and post-FMT samples. Successful FMTs were associated with a reduction in acylcarnitines, primary bile acids, along with an increase in secondary bile acids. We also identify microbial genes important for these changes, in particular carnitine metabolism genes encoded by many Enterobacteriaceae pre-FMT. Enterobacteriaceae also encoded many amino acid biosynthesis genes pre-FMT, potentially able to provide *C. difficile* with amino acids it is auxotrophic for. To further identify changes to the bile acid and amino acid pool, we performed multidimensional bile acid analyses by coupling liquid chromatography, ion mobility spectrometry, and mass spectrometry (LC-IMS-MS) separations. These measurements identified a shift from high microbial conjugated primary bile acids (AA-CA and AA-CDCA) pre-FMT to high secondary bile acids (AA-DCA) post-FMT. Finally, we investigate the role of BSHs in MCBA abundance and identify that a majority of BSHs post-FMT are encoded by the Lachnospiraceae Family. This study identifies specific changes in the microbial community and metabolic environment in response to a successful FMT, while providing insight into potential mechanisms that will help shape the development of future LBPs.

## Results

### Patient sample characteristics and clinical outcomes of FMT for rCDI

Samples were collected from fifteen patients enrolled for FMTs performed between January and December 2017, and processed between February 2017 and February 2018. Stool samples were collected pre-FMT, and post-FMT at time points of 2 weeks, 2 months, and 6 months (Figure 1A). Patient characteristics are described in Table 1 with 14 of the 15 patients (93.3%) clinically resolving their CDI after a single FMT. However, one subject, identified as recipient 9 (R9), developed rCDI three days after the initial FMT, and required an additional FMT. Thus, while the number of patients was 15, the total number of FMTs in this study was 16. Additionally, no patients had inflammatory bowel disease (IBD). One patient, R13, had significant immune suppression from recent chemotherapy for systemic amyloidosis. All FMTs studied were colonoscopic, with 14 instillations into the terminal ileum (TI) and 2 in the cecum. The failed FMT was one of the cecal infusions, and the successful second FMT was instilled into the TI, but also from a different donor. The longitudinal sampling timepoints for this patient are based off the successful FMT. Three other subjects had some persistence of diarrhea post-FMT, but throughout the course of their evaluations, rCDI was not found to be the cause. Pre-FMT stool samples were available for all 16 FMTs. Only 11 recipients submitted samples at 2 weeks, 10 recipients submitted samples at 2 months, and 8 recipients submitted samples at 6 months post-FMT. Two recipients, R11 and R15, did not submit any post-FMT samples. Stool samples from all recipients were analyzed using a multi-omic approach as shown in Figure 1A. Shallow shotgun sequencing by Diversigen identified 447 microbial species and 1,060,171 microbial genes. Untargeted metabolomic analysis by Metabolon identified 924 unique metabolites, while lipidomics using LC-IMS-MS identified 130 unique bile acids and MCBAs in the stool samples (Figure 1A, Table S8).

**Figure 1.**
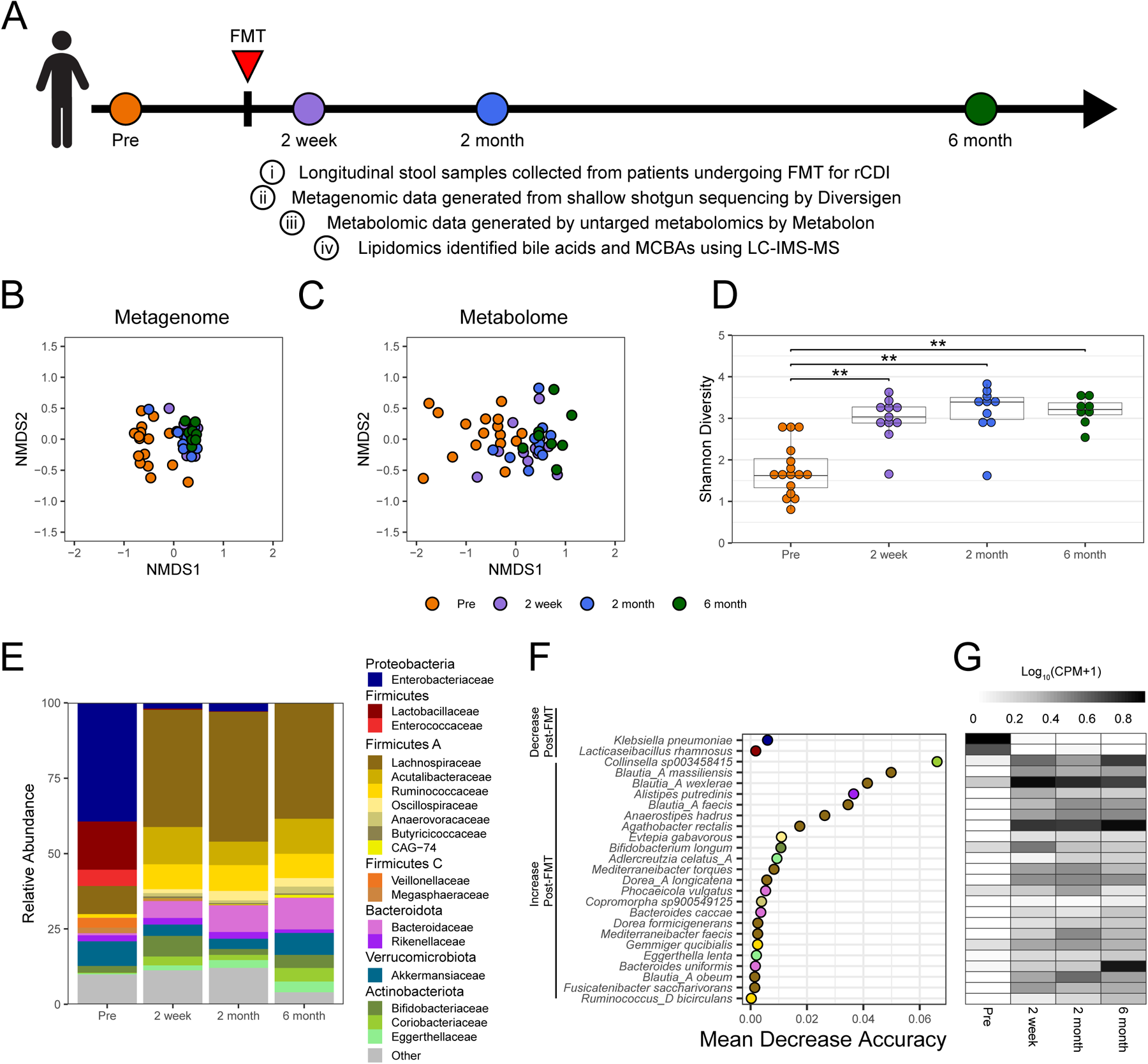
FMT significantly alters the structure and function of the gut microbiota. (**A**) Schematic of FMT stool sample collection from patients undergoing FMT for rCDI. Fecal samples were collected pre-FMT (n=16: orange), and then 2 week (n=11: purple), 2 month (n=10: blue), and 6 month (n=8: green) post-FMT for a total of (n=29) post-FMT samples. NMDS of Bray-Curtis dissimilarity of (**B**) bacterial species (stress=0.182) and (**C**) metabolites (stress=0.128) identified in stool samples from FMT patients. (**D**) Shannon Diversity of bacterial species identified in stool samples. Asterisks denote significance (** p ≤ 0.01) by pairwise Wilcoxon signed rank tests with Holm correction. (**E**) Average relative abundance of bacterial Family membership for each sampling timepoint in pre- and post-FMT samples. (**F**) Random Forest Analysis of species identified through metagenomics of stool samples The OOB error rate for RFA in determining pre- vs post-FMT is 4.44%. (**G**) Mean of log 10 transformation of counts per million (CPM) + 1 of reads assigned to each species as labelled in F.

**Table 1.** Characteristics of patients with recurrent *C. difficile* infection who underwent fecal microbiota transplantation, including 14 patients with initially successful FMT, and 1 with initially unsuccessful FMT.

|  | Successful FMT, N (%) | Unsuccessful FMT, N=1 |
| --- | --- | --- |
| Male sex | 5 (35%) | 0 |
| Median age* | 61.1 ($\pm 15.2$) | 63 |
| Caucasian race | 13 (86%) | 1 |
| Average BMI* | 25.3 ($\pm 3.9$) | 28.4 |
| Active smoker | 4 (31%) | 0 |
| Any alcohol use | 8 (33%) | 1 |
| Average number of *C. difficile* episodes (prior to FMT) | 4.5 (+/- 2.3) | 3 (NA) |
| Average number of *C. difficile* -related hospitalizations (prior to FMT) | 1.3 (+/- 1.3) | 0 |
| Severity of *C. difficile* -related diarrhea |  |  |
| 0-3 BMs daily |  |  |
| 3-5 BMs daily | 1 (7%) | 0 |
| >6 BMs daily | 4 (29%) | 1 |
|  | 9 (64%) | 0 |
| Active *C. difficile* infection (on suppressive antibiotics) at time of FMT | 9 (82%) | 1 |
| Average number of antibiotic courses for *C. difficile* in past year | 4.0 (+/- 1.6) | 2 |
| Prior therapies for *C. difficile* |  |  |
| Metronidazole | 9 (64%) | 0 |
| Vancomycin short course | 7 (50%) | 1 |
| Vancomycin taper | 13 (93%) | 1 |
| Fidaxomicin | 3 (21%) | 0 |
| Probiotic | 4 (29%) | 1 |
| Acid suppression (PPI or H2-blocker) |  |  |
| During previous infection only |  |  |
| During current infection | 4 (33%) | 0 |
|  | 5 (42%) | 0 |
| Median number of weeks after initial diagnosis that underwent FMT | 34.2 (20-138) | 21.4 |
| Endoscopic findings |  |  |
| Normal | 12 (86%) | 1 |
| Colitis, edema or other inflammatory change | 2 (14%) | 0 |
| Antibiotics within 2 weeks of FMT |  |  |
| Yes, vancomycin |  |  |
| Yes, non-vancomycin | 5 (63%) | 1 |
| None | 1 (13%) |  |
|  | 2 (25%) |  |
| Non- *C. difficile* antibiotic exposure after FMT | 5 (36%)† | 0 |
| Acid suppression post-FMT | 5 (36%) | 0 |

BM, bowel movement; FMT, fecal microbiota transplantation; IBD, inflammatory bowel disease; PPI, proton pump inhibitor.
If not all participants had a response recorded, the percentage was calculated from responders only
\*Where averages are presented, parentheses contain standard deviations, and medians contain ranges.
†Trimethoprim-Sulfamethoxazole, nitrofurantoin, doxycycline, amoxicillin, and unknown pneumonia antibiotic

### FMT significantly alters the structure and function of the gut microbiota

Significant differences were observed in the microbiome and metabolome between the pre- and post-FMT samples. Notably, shifts in the Bray-Curtis dissimilarity among the microbial species identified from metagenomics (p ≤ 0.001 by pairwise Adonis; Figure 1B), as well as the metabolome between pre- and all post-FMT time points were observed (p ≤ 0.001 by pairwise Adonis; Figure 1C). However, no significant differences in Bray-Curtis dissimilarity between post-FMT time points (2 week, 2 month, and 6 month) across any of these datasets. Additionally, the Shannon diversity of microbial species in stool samples was higher in all post-FMT samples compared to pre-FMT samples (p ≤ 0.01 by Wilcoxon signed rank test with Holm correction; Figure 1D). At the Phylum level, this corresponds to a decrease in Proteobacteria and Firmicutes, and an increase in Firmicutes A, Bacteroidota, and Actinobacteroidota post-FMT. FMT also corresponds with decreases in Family membership of Enterobacteriaceae (39.3% to <2.8%), Lactobacillaceae (16% to <0.5%), Enterococcaceae (5.4% to <0.05%), Veillonellaceae (3.2% to 0%), and Megasphaeraceae (1.8% to <0.8%) and increases in Lachnospiraceae (9.2% to 43.1%), Acutalibacteraceae(0.15% to 12.4%), Ruminococcaceae (1.1% to 8.6%), Oscillospiraceae (0.01% to 3.2%), Anaerovoracaceae (0.5% to 2.2%), Butyricicoccaceae (0% to 0.7%), CAG-74 (0% to 1%), Bacteroidaceae (0.7% to 10.5%), Rikenellaceae (2% to 2.2%), Bifidobacteriaceae (2.3% to 6.9%), Coriobacteriaceae (0.2% to 4.5%), and Eggerthellaceae (0.38% to 3.5%) (q ≤ 0.05 linear mixed models with Bonferroni correction; Figure 1E and Table S1). However, there was some variation between individuals within the same sampling timepoint (Figure S1). To further identify the bacteria at the species level that were driving the changes resulting from FMT, we used Random Forest Analysis (RFA), a machine learning algorithm. The Mean Decrease in Accuracy (MDA) reflects the importance of each species in determining whether a sample was correctly classified as pre- or post-FMT (Figure 1F). Of the 25 species identified as important (MDA >0), *Klebsiella pneumoniae* (Enterobacteriaceae) and *Lacticaseibacillus rhamnosus* (Lactobacillaceae) dominated the stool pre-FMT, and among the most important species post-FMT were *Collinsella sp003458415* (Coriobacteriaceae*), Blautia massiliensis*, *Blautia wexlerae*, *Blautia faecis, Anaerostipes hadrus,* and *Agathobacter rectalis* (Lachonospiraceae), and *Alistipes putrednis* (Rikenellaceae) (Figure 1F-G).

### Lipid metabolism is significantly altered between pre- and post-FMT

An untargeted metabolomics platform identified 924 metabolites of which 182 were significantly different between pre- and post-FMT (q ≤ 0.05 by linear mixed models with Bonferroni correction; Table S2). Specifically, 110 metabolites decreased in abundance, whereas 72 increased post-FMT (Table S3). RFA found 374 metabolites to be important (MDA >0), 220 of which decreased in abundance, and the other 154 increased in abundance post-FMT. We focused our analysis on the top 25 metabolites from each group that decreased or increased post-FMT, totaling 50 important metabolites (q ≤ 0.05 by linear mixed models with Bonferroni correction; Figure 2A, Table S2) and their abundances are illustrated in Figure 2B.

**Figure 2.**
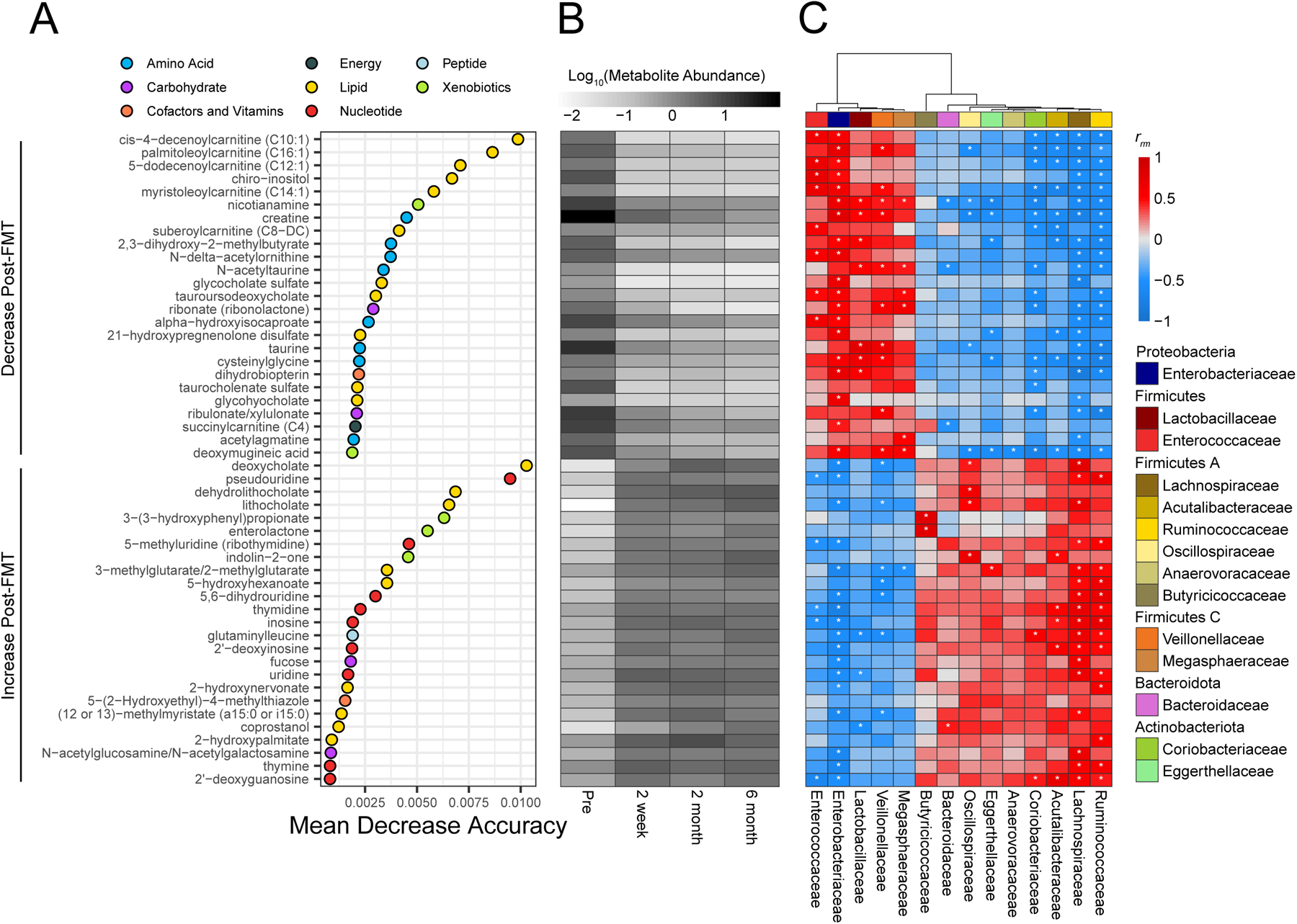
Lipids: Acylcarnitines and bile acids are the most predictive metabolites able to distinguish between pre- and post-FMT samples. (**A**) RFA of metabolites defined by untargeted metabolomics in stool samples. Metabolites were categorized into those that decrease after FMT or increase after FMT and the top 25 metabolites from each category are plotted. The OOB error rate for RFA in determining pre- vs post-FMT is 8.89%. (**B**) Mean of log 10 transformed, median scaled, minimum imputed abundance of important metabolites identified by RFA for each time point. (**C**) Repeated measure correlation between abundance of important metabolites identified by RFA and relative abundance of microbial Families that were significantly different between pre- and post-FMT. Significant repeated measure correlations (p ≤ 0.05) by linear model with Benjamini and Hochberg correction are marked with an asterisk.

Lipids emerged as a key distinguishing factor between pre- and post-FMT, with 20 out of the top 50 metabolites belonging to this super pathway category designated by Metabolon (Figure 2A-B). The largest classes of lipids that decreased post-FMT were acyl-carnitines and bile acids, which were also among the most important metabolites as denoted by the RFA. Specifically, six acylcarnitines, cis-4-decenoylcarnitine, palmitoleoylcarnitine, 5- dodecenoylcarnitine, myristoleoylcarnitine, suberoylcarnitine, and succinylcarnitine were among the most important metabolites by MDA that decreased in abundance post-FMT. Four conjugated bile acids decreased post-FMT (glycocholate sulfate, tauroursodeoxycholate (TUDCA), taurocholenate sulfate, and glycohyocholate (GHCA)), whereas three unconjugated secondary bile acids increased post-FMT including deoxycholate (DCA), dehydrolithocholate (3- oxo LCA), and lithocholate (LCA). Amino acids were among the most important by RFA that decreased post-FMT (Figure 2A-B). In contrast, nucleotides were among the top metabolites that increased post-FMT (Figure 2A-B). Carbohydrates, cofactors and vitamins, energy, peptides, and xenobiotics were also observed, but to a lesser degree and importance based on the MDA score.

To better define the relationship between the microbiota and the metabolome, while accounting for multiple post-FMT timepoints for each recipient, we performed repeated measure correlations (rmcorr) between metabolites and the relative abundance of bacterial Families that were significantly different between pre- and post-FMT (Figure 2C)^47^. The acylcarnitines that were high in pre-FMT and decreased post-FMT positively correlated with Enterobacteriaceae, Enterococcaceae, and Veillonellaceae. The bile acids that were highest in pre-FMT (glychocholate sulfate, TUDCA, taurochenolate sulfate, and GHCA) significantly positively correlated with Enterobacteriaceae, Enterococcaceae, and Megasphaeraceae (p ≤ 0.05 by linear model with Benjamini and Hochberg correction; Figure 2C). The unconjugated secondary bile acids that increased post-FMT (DCA, 3-oxo LCA, and LCA) positively correlated with Lachnospiraceae and Oscillospiraceae, while the nucleotides that increased post-FMT correlated with Lachnospiraceae, Ruminococcaceae, Acutalibacteracaeae, and Coriobacteriaceae (p ≤ 0.05 by linear model with Benjamini and Hochberg correction; Figure 2C). These findings suggest an association between high acylcarnitine abundances pre-FMT and Enterobacteriaceae, and high secondary bile acid abundances post-FMT and Lachnospiraceae.

### Enterobacteriaceae encode the majority of carnitine metabolism genes that are dominant pre-FMT

To further identify relationships between the microbiota and lipidome, we evaluated microbial metabolic pathways associated with changes in lipid abundance. Carnitine metabolism emerged as one of these pathways. Coinciding with the decrease in acylcarnitines, there was also a decrease in carnitine metabolism at every time point post-FMT (p ≤ 0.01 by Wilcoxon signed rank test with Holm correction; Figure 3A). Furthermore, we identified carnitine metabolic pathways across 11 species belonging entirely to the Enterobacteriaceae Family, which positively correlated with acylcarnitine abundance (Figure 2C, Figure 3B). To further investigate the relationship between carnitine metabolism and acylcarnitines, we performed rmcorr between acylcarnitines, alongside carnitine, and the carnitine metabolic pathways of each species. Untargeted metabolomics identified 30 acylcarnitines, 11 of which were significantly different between pre- and post-FMT (p ≤ 0.05 by linear mixed models with Bonferroni correction; Table S2). The abundance of the carnitine metabolic pathway was encoded by seven species; *Salmonella enterica, Enterobacter cloacae, Escherichia fergunsonii, Citrobacter youngae, Salmonella bongori, Citrobacter portucalensis*, and *Proteus mirabilis* and positively correlated with acylcarnitines (p ≤ 0.05 by linear model with Benjamini and Hochberg correction; Figure 3C). Only *P. mirabilis*’ carnitine metabolic pathway had a significant correlation with carnitine itself.

**Figure 3.**
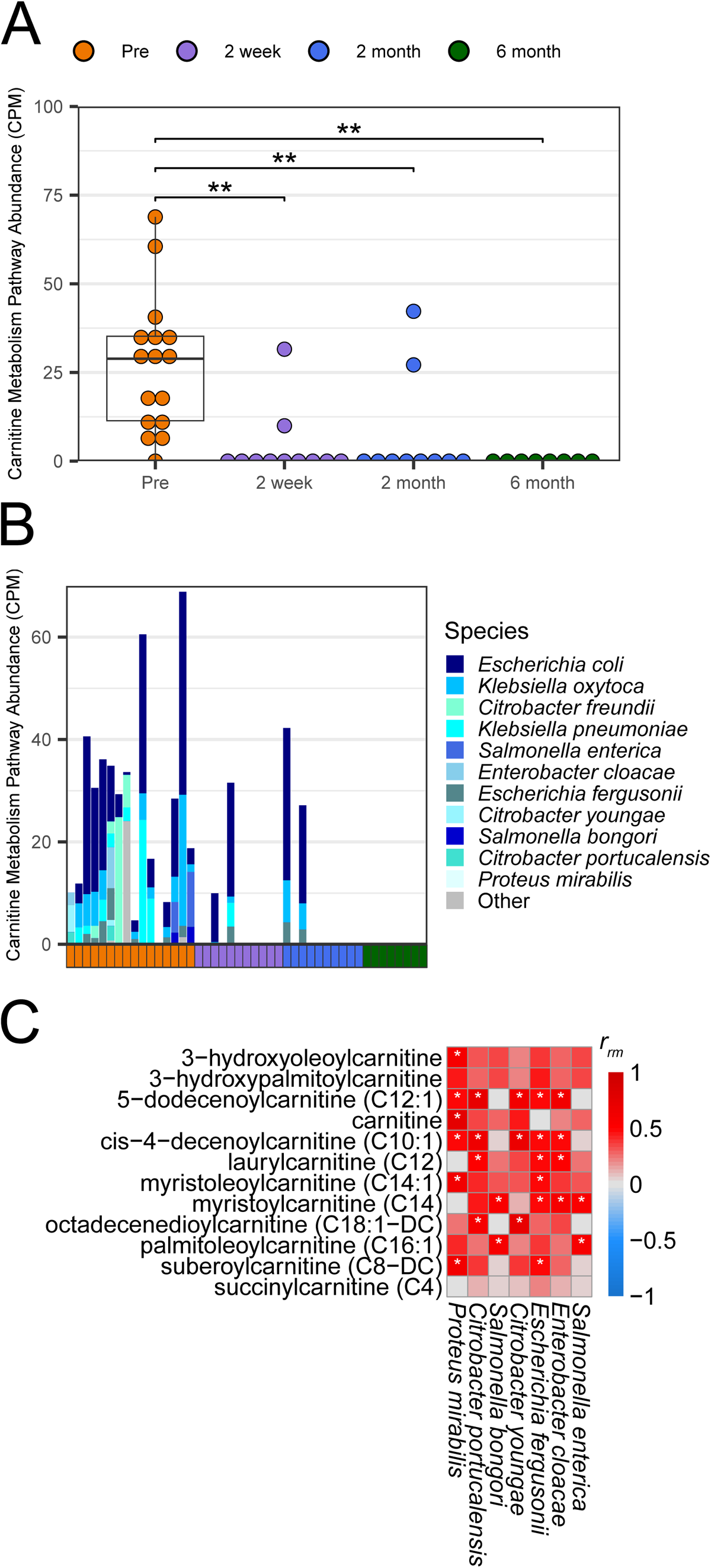
Pre-FMT samples are dominated by Enterobacteriaceae which encode the majority of carnitine metabolism genes. (**A**) Counts per million (CPM) of genes belonging to the carnitine metabolism UniPathway identified via shallow shotgun metagenomics. Asterisks denote significance (** p ≤ 0.01) by pairwise Wilcoxon signed rank tests with Holm correction. (**B**) Stacked bar plot of CPM carnitine metabolism UniPathway split by the species that encode it for each stool sample. Genes categorized as “Other” were identified via BLASTx. (**C**) Repeated measure correlation between acylcarnitines determined to be significantly different between pre- and post-FMT by linear mixed models with Bonferroni correction (q ≤ 0.05) and species-specific carnitine metabolism genes. Significant correlations (p ≤ 0.05) by linear model with Benjamini and Hochberg correction are marked with an asterisk.

### Stickland fermentation products and biosynthesis of amino acids *C. difficile* is auxotrophic for decrease post-FMT

Amino acids are essential for *C. difficile* growth, important for energy production using Stickland fermentation, and able to shape the gut microbiota due to their auxotrophies ^32,48^. Therefore, we focused our analysis on the amino acids and dipeptides identified in untargeted metabolomics. RFA identified 25 important amino acids or dipeptides that decreased and 18 that increased post-FMT (Figure S2). There were two Stickland products that significantly decreased post-FMT, alpha-hydroxyisocaproate and 5-aminovalerate which result from reductive leucine and reductive proline Stickland fermentation, respectively ^49,50^. These two Stickland products correlated with Enterobacteriaceae, Lactobacillaceae, Enterococcaceae, Veillonellaceae, and Megasphaeraceae (p ≤ 0.05 by linear model with Benjamini and Hochberg correction; Figure S2). We also noted higher N-acetylated amino acids pre-FMT, indicating peptide cleavage, and higher abundance of dipeptides post-FMT (Figure S2).

Since *C. difficile* is auxotrophic for six amino acids: cysteine, isoleucine, leucine, proline, tryptophan, and valine, it requires other members of the microbiota or its environment to provide them. Therefore, we aimed to identify which microbes in pre-FMT samples encode genes for amino acid biosynthesis (Figure 4). Enterobacteriaceae was a major contributor of amino acid biosynthesis pre-FMT and its members synthesize five of the six amino acids *C. difficile* is auxotrophic for including cysteine, isoleucine, leucine, proline, and tryptophan (Figure 4). For leucine, proline, and tryptophan there is either no reduction or a moderate reduction in the amount of these biosynthesis pathways pre- and post-FMT. However, a notable shift occurs in the taxa for these functions (Figure 4). Bacteroidaceae also contributes to the biosynthesis of other amino acids such as arginine, homocysteine, and lysine, while Bifidobacteria contribute to the majority of histidine post-FMT and the *C. difficile* auxotrophy leucine. No taurine or valine biosynthesis pathways were identified (Figure 4). Furthermore, amino acid biosynthesis pathways comprise a large portion of the important pathways by MDA when comparing pre- and post-FMT (Figure S3).

**Figure 4.**
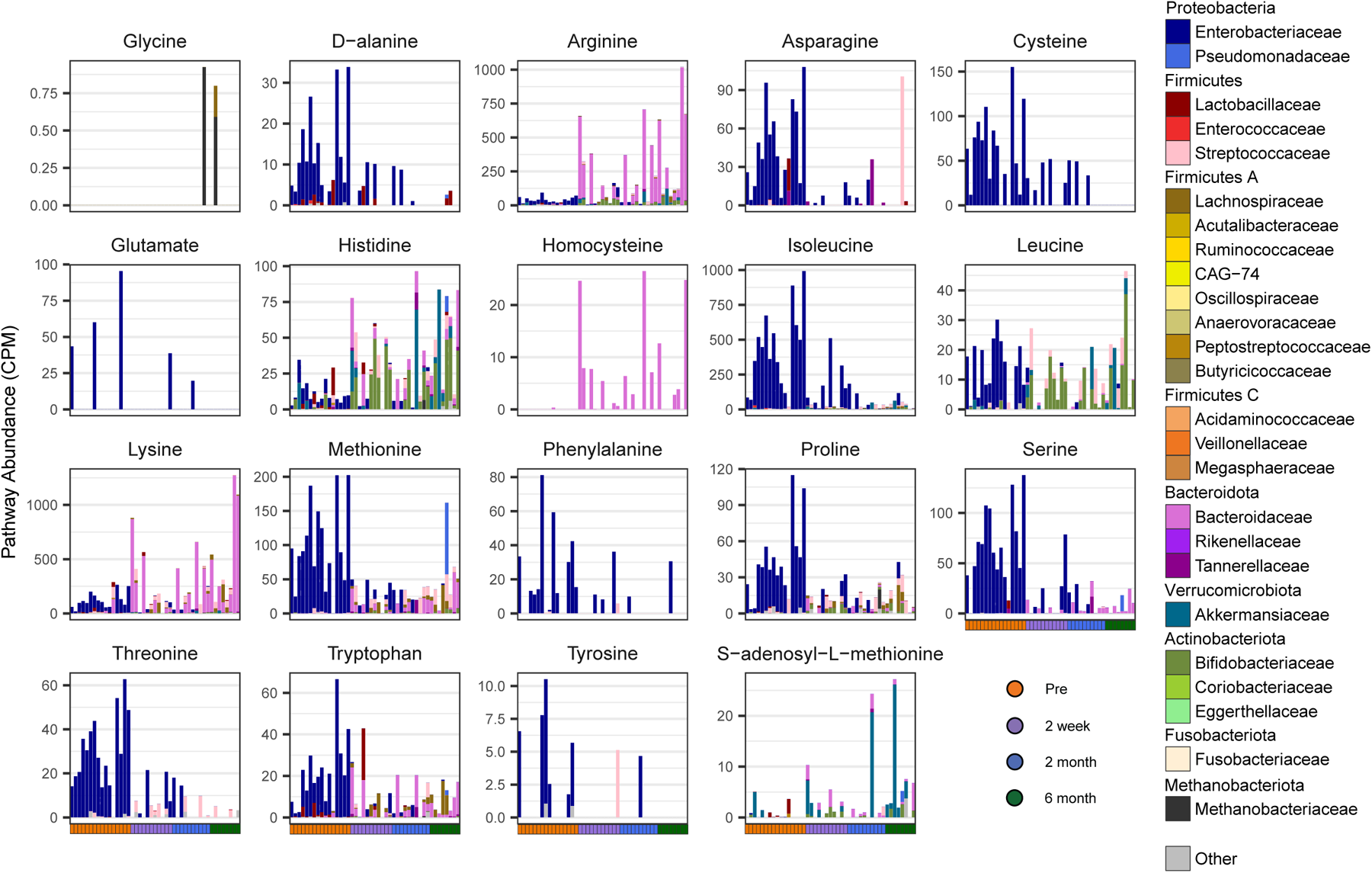
Pre-FMT samples have an increase in microbial biosynthesis of amino acids *C. difficile* is auxotrophic for. Stacked barplots of counts per million (CPM) of genes belonging to the labeled amino acid biosynthesis UniPathway, split by the family that encodes it. Each bar represents one patient timepoint. Genes categorized as “Other” were identified via BLASTx.

### Microbial conjugation of secondary bile acids dominates post-FMT samples

To further characterize the relationship between bile acids, amino acids, and FMT, we performed specific extractions and lipidomic analyses to target bile acids and MCBAs in the stool samples collected pre- and post-FMT ^45^. A targeted library containing 208 unique bile acids was utilized to evaluate the LC-IMS-MS measurements. Of the 208 bile acids, 130 were detected in at least one sample and 43 had significantly different abundances between the pre- and post-FMT samples (q ≤ 0.05 by linear mixed models with Bonferroni correction; Table S4). Subsequent RFA identified 88 bile acids as important including all 43 significantly different bile acids (Figure 5A).

**Figure 5.**
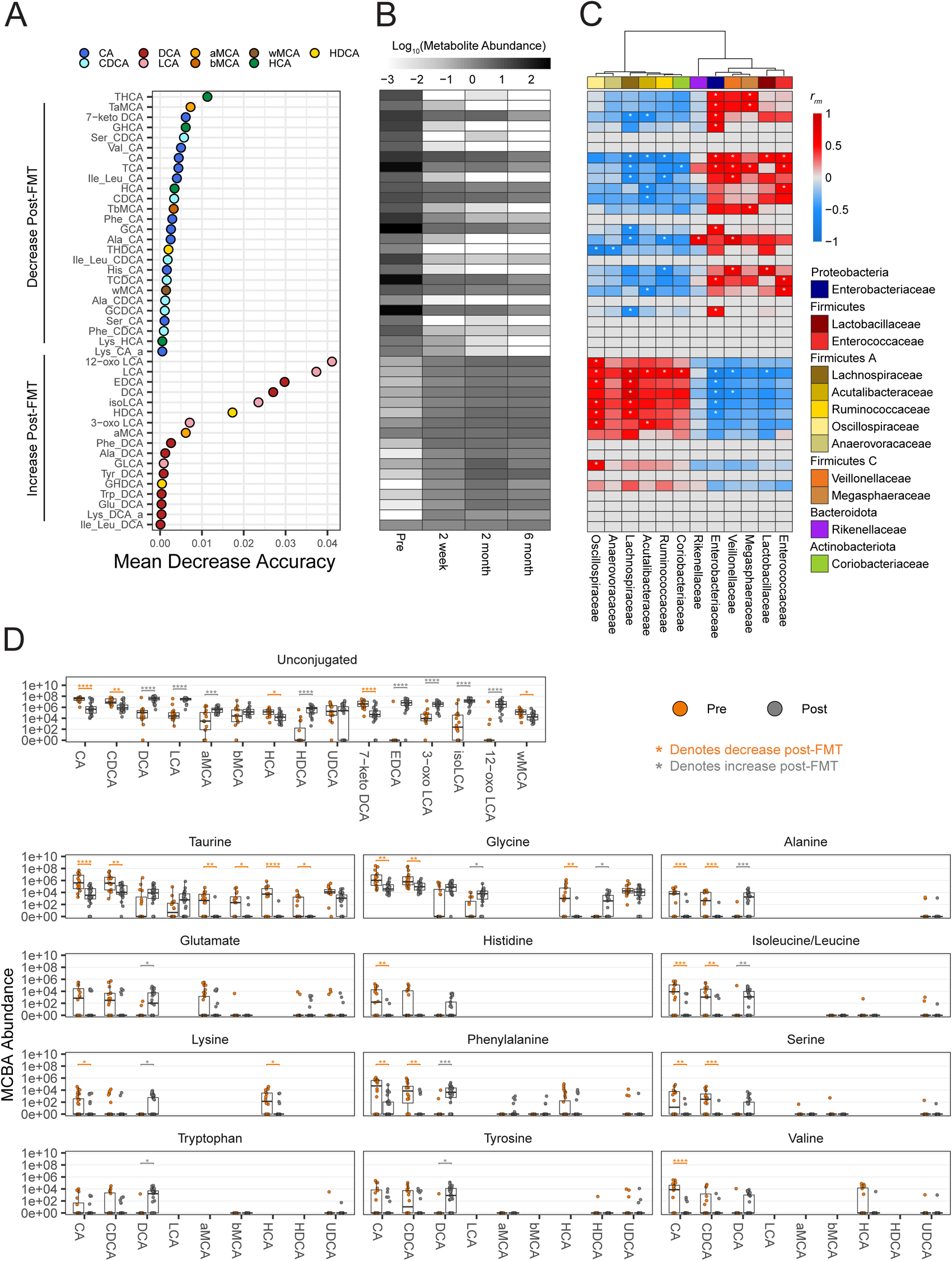
Microbially conjugated bile acid pool post-FMT favors secondaray bile acids. (**A**) RFA of MCBAs identified by targeted metabolomics in stool samples. The OOB error rate for RFA in determining pre- vs post-FMT is 6.67%. (**B**) Mean of log 10 transformed, median scaled abundance of important metabolites identified by random forest analysis for each time point. (**C**) Repeated measures correlation between abundance of important MCBAs identified by RFA and relative abundance of microbial families that were significantly different between pre- and post-FMT. Significant repeated measure correlations (p ≤ 0.05) by linear model with Benjamini and Hochberg correction are marked with an asterisk. (**D**) MCBA abundance for each amino acid/sterol core combination found to be significant between timepoints or after FMT by linear mixed models. Asterisks denote significance (* q ≤ 0.05, ** q ≤ 0.01, *** q ≤ 0.001, **** q ≤ 0.0001) by linear mixed models.

Of these 43 statistically significant bile acids, 26 decreased post-FMT, whereas the other 17 increased (Figure 5A-B). Predominantly, primary bile acid amidates decreased post-FMT, with nine containing cholate (CA) sterol cores and six containing chenodeoxycholate (CDCA) (Figure 5A-B). In contrast, secondary bile acid amidates were observed to increase post-FMT with six having a DCA sterol core (Figure 5A-B). Similar to the results in Figure 2, DCA, 3-oxo LCA, and LCA were among the most important bile acids to return post-FMT. However, increases in 12-oxolithocholate (12-oxoLCA), 3-epideoxycholate (EDCA), isolithocholate (isoLCA) and hyodeoxycholate (HDCA) were also significant using this targeted approach. These findings reflect the importance of LCA and its derivatives in post-FMT recovery. While our bile acid library lacks MCBAs with LCA derived sterol cores due to the absence of synthesized standards, the presence of MCBAs with other secondary bile acid cores, specifically DCA, were found to be highly important post-FMT.

Correlating this targeted panel of bile acids with the microbiome, we also discovered a significant association with bacterial Families. Isoleucine/leucine, alanine, and histidine conjugated CA decreased in abundance post-FMT and positively correlated with one or more of Lactobacillaceae, Veillonellaceae, and Rikenellaceae (p ≤ 0.05 by linear model with Benjamini and Hochberg correction; Figure 5C). Families including Enterobacteriaceae, Enterococcaceae, Veillonellaceae, and Megasphaeraceae positively correlated with at least one traditional host associated (taurine or glycine) primary bile acid amidates that decreased post-FMT. However, there is also the potential that these amino acids are being conjugated by the microbiota. Additionally, Lachnospiraceae, Acutalibacteraceae, Ruminococcaceae, Oscillospiraceae, and Coriobacteriaceae significantly positively correlated with unconjugated secondary bile acids (12- oxo LCA, LCA, EDCA, DCA, isoLCA, HDCA, and 3-oxo LCA) that increased post-FMT (p ≤ 0.05 by linear model with Benjamini and Hochberg correction; Figure 5C). Oscillospiraceae, in particular, showed a positive correlation with GLCA, which may potentially have been host-conjugated. Nonetheless, no other Family showed significant positive correlations with secondary bile acid amidates. These finding suggest Lactobacillaceae, Veillonellaceae, and Rikenellaceae may be driving MCBA production pre-FMT, while Lachnospiraceae, Acutalibacteraceae, Ruminococcaceae, Oscillospiraceae, and Coriobacteriaceae impact the secondary bile acid pool post-FMT.

Significant alterations in the abundance of multiple unconjugated bile acids were also observed between pre- and post-FMT. Specifically, CA, CDCA, HCA, 7-keto DCA, and ωMCA decreased in abundance post-FMT, while DCA, LCA, αMCA, HDCA, EDCA, 3-oxoLCA, isoLCA, and 12-oxoLCA increased in abundance post-FMT. Although some exceptions exist, this trend represents a decrease in primary bile acid amidates coupled with an increase in secondary bile acid amidates. Conjugation of glycine, alanine, isoleucine/leucine, and phenylalanine to primary bile acids CA and/or CDCA decreased post-FMT, while conjugation to the secondary bile acid DCA increased (p ≤ 0.05 by linear mixed models with Bonferroni correction; Figure 5D). A preference for glutamate, histidine, serine, tryptophan, tyrosine, and valine conjugation to specific sterol cores was indicated by either a decrease in primary bile acid amidates or increase in secondary bile acid amidates (Figure 5D). Although non-significant changes between pre- and post-FMT in other MCBAs were also observed (Figure S4), these findings highlight that both the bile acids and the amino acids that are being conjugated by the microbiota are impacted by FMT.

### Bile salt hydrolases increase post-FMT

Bile salt hydrolases (BSHs) are enzymes that have recently been shown to catalyze the conjugation of bile acids and are also regarded as gatekeepers of bile acid modifications, as they remove amino acids from the sterol core, allowing further modifications to occur ^43,44,46^. Therefore, we investigated BSHs presence in pre- and post-FMT stool samples to define the changes in the metagenome correlate with the metabolome. We identified 153 BSHs from 89 species (Figure S5), with an increase of BSH genes between pre-FMT and all post-FMT time points (p ≤ 0.05 by Wilcoxon signed rank test with Holm correction; Figure 6A). BSHs were further classified based on their specificity loop ^46^. There was an increase in glycine-preferring BSHs between pre-FMT and 2 week post-FMT (p ≤ 0.05 by Wilcoxon signed rank test with Holm correction; Figure 6B), many of which. These glycine-preferring BSHs also contain the X-S-R-X motif, implicating their roles in MCBAs activity ^51^. The BSHs that returned post-FMT were encoded largely by Firmicutes A and Bacteroidota, primarily from the Lachnospiraceae Family as well as some from Bacteroidaceae (Figure 6C). We did not detect any BSHs in one post-FMT sample, R3, at 2 months (Figure 6C). To further clarify which members of the microbiome contribute to the BSH pool, we summarized the total amount of BSH encoded by each species (Figure S5). There was a broad range of species encoding BSHs, though BSHs encoded by *Collinsella aerofaciens, Blautia obeum,* and *Blautia wexlerae* were among species that encoded the most BSHs across post-FMT samples. These species as well as other top BSH contributors were also among the most important species between pre- and post-FMT (Figure S5 and 1F).

**Figure 6.**
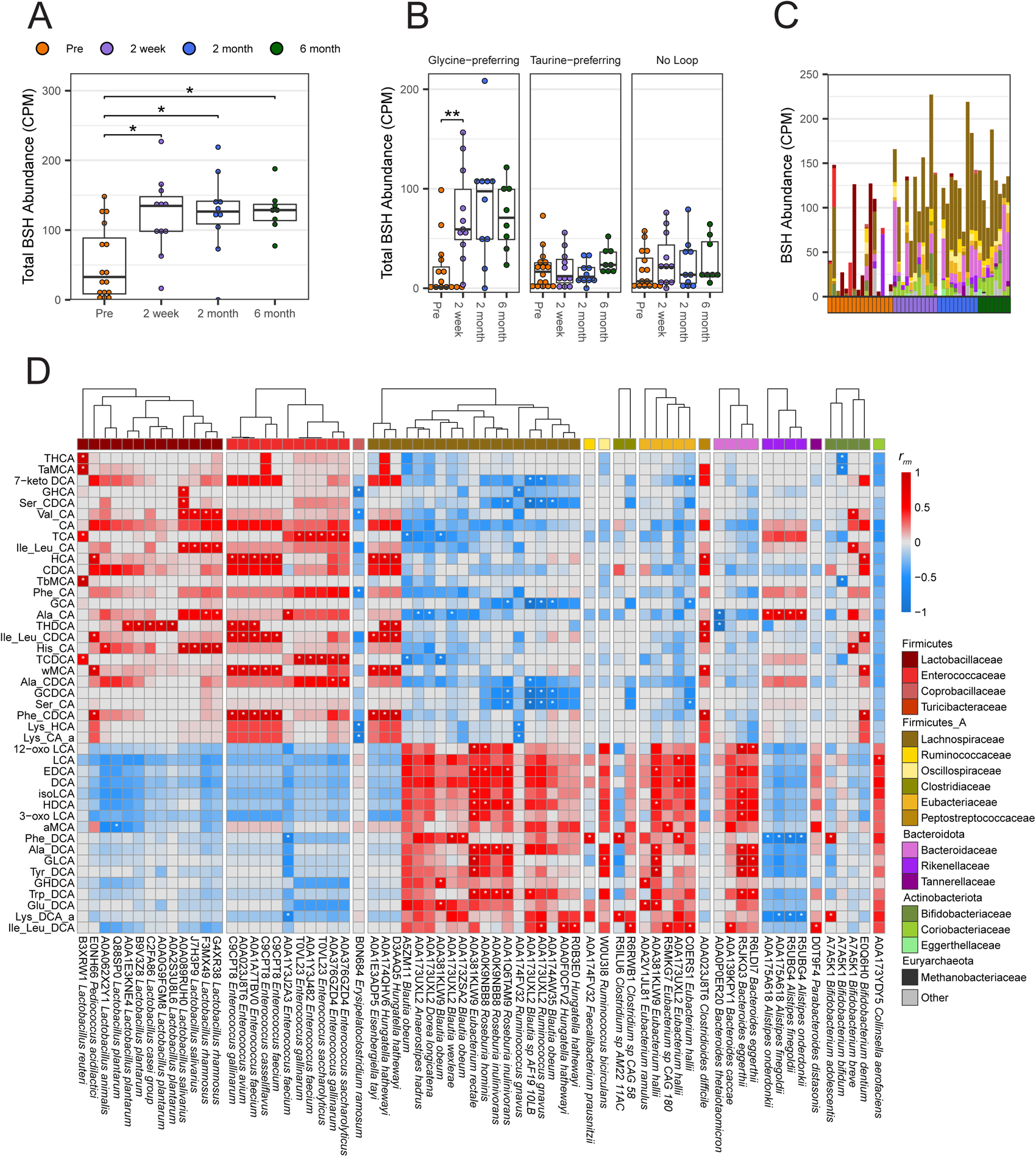
Bile salt hydrolases significantly increase post-FMT and are encoded mainly by the Lachnospiraceae Family. Genes were identified as BSHs if their UniRef90 protein ID contained “Bile Salt Hydrolase” or “Choloylglycine Hydrolase”. (**A**) Counts per million (CPM) of BSH genes identified via shallow shotgun metagenomics. (**B**) CPM of BSH genes separated by loop specificity determined by MUSCLE alignment of UniRef90 sequences. Asterisks within boxplots denote significance (* p ≤ 0.05, ** p ≤ 0.01) by pairwise Wilcoxon signed rank tests with Holm correction. (**C**) Stacked bar plot of BSH genes split by the Family that encode it for each stool sample. Genes categorized as “Other” were identified via BLASTx and do not have species information. (**D**) Repeated measure correlation between MCBAs identified by targeted metabolomics and observed to be significantly different between pre- and post-FMT and BSH genes identified by shallow shotgun sequencing. Significant correlations (p ≤ 0.05) by linear model with Benjamini and Hochberg correction are marked with an asterisk.

To further characterize these BSHs and their impact on the MCBA pool, we correlated each individual BSH with targeted bile acid abundances (Figure 6D). Generally, Lactobacillaceae and Enterococcaceae encoded BSHs positively correlated with the unconjugated or amidated primary bile acids that decreased post-FMT, while Lachnospiraceae, Eubacteriaceae, and Bacteroidaceae encoded BSHs positively correlated with the unconjugated or amidated secondary bile acids that increased post-FMT. This may reflect a role these BSHs play in the production of MCBAs in their respective environments. BSHs are labelled based on their UniProt90 identifiers, which groups them based on 90% protein identity. Moreover, certain groups of BSHs were observed multiple times; A0A173UXL2, A0A381KLW9, A7A5K1, and C9CPT8 were each observed at least three times with A0A173UXL2 being the most common. A0A173UXL2 and A0A381KLW9 were found most commonly among Lachnospiraceae and positively correlated with 12-oxo-LCA, EDCA, isoLCA, HDCA, 3-oxoLCA, Phe_DCA, Ala_DCA, GLCA, Tyr_DCA, GHDCA, Trp_DCA, Glu_DCA, and Ile_Leu_DCA. A7A5K1 was found in Bifidobacteriaceae and was correlated with Val_CA, Ile_Leu_CA, His_CA, Phe_DCA, and Lys_DCA. C9PT8 was found among Enterococcaceae and correlated generally with HCA, THDCA, Ile_Leu_CA, ωMCA, and Phe_CDCA. These findings highlight the specificity of BSHs for certain bile acids and identify groups of BSHs that may play a role in modifying the bile acid pool.

Following BSH-mediated deconjugation, additional bile acid altering enzymes can further modify the sterol core of bile acids, making secondary bile acids. There were shifts in the abundance of some of these enzymes as well, although many of them did not meet significance requirements (Figure S6). *baiA* in the *bai* operon was the only gene that was significantly different pre- vs post-FMT and was encoded primarily by Lachnospiraceae (p ≤ 0.05 by Wilcoxon signed rank tests with Holm correction; Figure S6). The *bai* operon is able to produce secondary bile acids and changes in the abundance of this gene is reflected in changes in abundance of secondary bile acid sterol cores (Figure 5D).

## Discussion

In this study we performed metagenomics, metabolomics and lipidomics using different platforms on the stool of patients undergoing FMT for rCDI to determine mechanisms of a successful FMT. Significant increases in alpha and beta diversity have previously been described after FMT for rCDI. Though our study adds additional resolution at the strain level.

We also noted, alongside other studies ^23,24,26^, a significant reduction in antimicrobial resistance genes (AMRs) (Table S5). Members of the Enterobacteriaceae Family dominated the pre-FMT environment and significantly decreased post-FMT, which gave way to members of the Lachnospiraceae Family, and an overall increase in bacterial diversity. Shifts in the microbiota were accompanied by large changes in the lipidome, specifically acylcarnitines and bile acids. Acylcarnitine levels were high pre-FMT, and there was a shift from conjugated primary bile acids to secondary bile acids from pre- to post-FMT. Acylcarnitines correlated with members from the Enterobacteriaceae Family, which encoded the majority of carnitine metabolism genes pre-FMT. Enterobacteriaceae also encoded many genes involved in the biosynthesis of amino acids that *C. difficile* is auxotrophic for. Stickland fermentation products were also higher pre-FMT when compared to post-FMT. MCBAs shifted from primary bile acid amidates pre-FMT to secondary bile acid amidates post-FMT. BSHs, enzymes able to deconjugate and reconjugate MCBAs, significantly increased post-FMT, and were encoded primarily by members of the Lachnospiraceae Family.

The pre-FMT stool environment is enriched in acylcarnitines. Acylcarnitines are important for long-chain fatty acid β-oxidation by the host ^52^. The release of acylcarnitines from within the cell is associated with inflammation mediated apoptosis and mitochondrial dysfunction in IBD ^53,54^. Enterobacteriaceae can grow on acylcarnitines *in vitro* indicating they may also possess a mechanism to liberate carnitine from acylcarnitines ^53^. Gut bacteria utilize carnitine for growth through use of the *cai* operon, which is found in a higher abundance in acylcarnitine rich environments ^55^. High levels of acylcarnitines are also found alongside high levels of Enterobacteriaceae in pediatric IBD patient samples ^53^. Carnitine metabolism genes in our study were encoded entirely by Enterobacteriaceae and positively correlated with the abundance of acylcarnitines. At this time, it is not known if *C. difficile* is able to grow on these lipids, as we were not able to find any annotated carnitine metabolism genes in *C. difficile*.

The Enterobacteriaceae in pre-FMT stool also encoded many genes important for the biosynthesis of *C. difficile* auxotrophic amino acids; cysteine, isoleucine, leucine, proline, tryptophan, and valine ^32^, and amino acids that other members of the gut microbiota are auxotrophic for including asparagine, methionine, phenylalanine, serine, threonine, and tyrosine ^48^. Amino acid auxotrophies in the gut shape diversity and stability of the microbiota ultimately impacting gut health ^48^. A recent study showed that stool from successful FMT donors was enriched in many amino acid biosynthesis genes, which suggests that cross-feeding of amino acids could be important for successful FMT outcomes ^29^. In another study *Clostridium sardiniense* was able to provide ornithine and other fermentable amino acids *C. difficile,* increasing pathogen burden and morbidity in gnotobiotic mouse model ^56^. Enterococci produces fermentable amino acids such as leucine and ornithine to *C. difficile*, thereby increasing its fitness in the gut ^57^. *C. difficile* also provides nutrients these organisms need as well, hence the term cross-feeding. *C. difficile* can also provide host heme through toxin mediated liberation to Enterococci in a mouse model of CDI ^57^. In this study we saw a significant decrease in the abundance of Enterococcaceae post-FMT and we did not observe any amino acid biosynthesis attributed to Enterococcaceae. All of the amino acid biosynthesis genes identified in this study were attributed to Enterobacteriaceae, which were also highest pre-FMT. Competition for key amino acids could also play a role in restoring colonization resistance against *C. difficile*. In a gnotobiotic mouse model of CDI, competition for Stickland metabolites between commensal Firmicutes (*Clostridium hiranonis, Clostridium leptum, Clostridium scindens*) and *C. difficile* was enough to prevent weight loss from CDI ^58^. There was some correlative evidence in our study that the Butyricicocaceae and Lachnospiraceae may be competing for amino acids in the post-FMT gut, although full metabolic modelling was limited by shallow sequencing.

A major focus of this study was to look at the contribution of not only amino acids, but bile acids, specifically newly defined MCBAs and the BSHs that drive them. A majority of BSHs in post-FMT stool were encoded by members of the Lachnospiraceae, a Family normally highlighted for its role in secondary bile acid production via the *bai* operon. Though we did also observe the presence of genes involved in secondary bile acid metabolism, such as *baiA*, also encoded by Lachnospiraceae alongside significant increases in the amount of many unconjugated secondary bile acids, which are known to inhibit multiple stages of the *C. difficile* life cycle ^37,59^. The Lachnospiraceae encoded BSHs also disproportionately contained the selectivity loop motif that has been implicated in dictating BSH preference for aromatic MCBAs, specifically primary bile acid amidates ^51^. Primary bile acid amidates generated by BSHs encoded by Lactobacilli strains have been shown to reduce *C. difficile* germination, vegetative growth, and toxin expression ^46^. Lactobacillaceae encoded BSHs correlated with primary bile acid amidates in our study, alongside Enterococcaceae encoded BSHs, and were highest in abundance in pre-FMT stool. Lachnospiraceae, Eubacteriaceae, and Bacteroidaceae encoded BSHs correlated with secondary bile acid amidates and were highest in post-FMT stool. The exact role these secondary bile acid amidates play in CDI and successful FMT has yet to be determined. This is in contrast to what has been observed in IBD, where primary bile acid amidates are higher in abundance in Crohn’s disease compared to healthy individuals, while there were no changes in secondary bile acid amidates ^60^. In patients with Crohn’s disease, secondary bile acid amidates are lower in symptomatic compared to asymptomatic patients ^55^.

This study used a comprehensive suite of omics techniques to define a successful FMT for rCDI, however there were some limitations to the study. First, we used shallow shotgun sequencing samples with a wide range of alpha diversity. We expect that samples that were >50% of a single species had more sequencing depth for that species relative to other more diverse samples and therefore may have had more complete metabolic pathways identified. Due to the conflicts between databases in Humann 3 and Metaphlan 4 some genes may have been classified as belonging to a different organism previously grouped in within the same pangenome. Since the FMT induced large shifts in both the microbiome and metabolome, we acknowledge some correlations may reflect these changes more so than specific associations between microbes and metabolites. Additionally, in our lipidomic analyses even with the advanced LC-IMS-MS platform, some of the MCBA isomers still exhibit overlap. For example, the similarity in retention time and collision cross section (CCS) for Leu-BA and Ile-BA in our synthetic library poses challenges for their definitive assignments ^45^. The scope of our library was also limited due to the selective utilization of bile acids in the synthesis process, a factor that is critical for the identification of MCBAs, given that it depends exclusively on the range of synthesized compounds available. Currently, our library includes MCBAs created by eight bile acids (CA, CDCA, DCA, HDCA, UDCA, αMCA, βMCA, and γMCA) conjugated with 22 amino acids ^61^. Looking forward, we anticipate expanding this library by incorporating a broader spectrum of MCBAs with different bile acid cores due to the synthesis of new standards. We note that patient sex had an impact on a small number of metabolites and metagenomic data, though our study was not appropriately powered to definitively make specific claims regarding sex (Table S6). Finally, though steps were taken to ensure normality for mixed effect linear models, that assumption was likely not met for every one of the thousands of measurements from metagenomics, metabolomics and lipidomics data.

The relationship between high acylcarnitines pre-FMT and *C. difficile* needs to be explored as this could be a potential mechanism to encourage Enterobacteriaceae growth and production of amino acids to further feed *C. difficile*. While other species have been observed to cross-feed with *C. difficile,* Enterobacteriaceae has not. Determining the specific relationship between the gut microbiota and MCBAs, particularly the source of amino acids that are conjugated and if conjugated amino acids can be used in metabolism will be important. In addition, confirming the impact of these secondary bile acid amidates on the *C. difficile* life cycle will further inform the role of these MCBAs and their importance during *C. difficile* colonization and infection. Lachnospiraceae were also identified as important contributors to the BSH pool and potentially MCBA production, in addition to 7α-dehydroxylation. Not much is known about BSHs encoded by this Family, despite their contribution to secondary bile acid metabolism. Uncovering the role they play in modifying the bile acid pool and sequestering amino acids will help guide research on MCBAs and LBPs. These findings will likely also apply to other diseases, particularly IBD. Overall, lipids were the largest class of metabolites significantly different and important by RFA pre- vs post-FMT. While our study highlights acylcarnitines and bile acids, more research is needed to determine how other types of lipids impact *C. difficile* and other intestinal diseases.

### Significance

Recurrent *C. difficile* infection (rCDI) is an urgent public health threat for which the last resort and lifesaving treatment is a fecal microbiota transplant (FMT). New Live Biotherapeutic Product s (LBPs) or microbiota-focused therapies (Rebyota and Vowst) were just approved by FDA in 2023 as a move to make FMTs safer and more homogeneous. However, the exact mechanisms which mediate a successful FMT are still not well understood. We define changes to the microbiome, metabolome, and lipidome of stool samples acquired from patients undergoing FMT for rCDI. We found high acyl-carnitines pre-FMT, an environment indicative of host cell lysis and ideal for Enterobacteriaceae growth. These Enterobacteriaceae were also observed to encode all of the carnitine metabolism in the gut and have the ability to synthesize amino acids that *C. difficile* is auxotrophic for. Bile salt hydrolases (BSHs) and other bile acid altering enzymes, particularly those encoded by Lachnospiraceae, increased in abundance after FMT. These changes were associated with changes to the bile acid pool, including changes in abundance of MCBAs. Specific BSHs correlated with the abundance of certain MCBAs, indicating BSH mediated generation of MCBAs is specific. These changes were observed alongside increases in secondary bile acids after FMT known to inhibit *C. difficile*. Our results provide insights into how the microbiome shapes the metabolome and lipidome in response to FMT. This information is critical for tailoring more targeted LBPs for the treatment of rCDI and other intestinal diseases.

## Acknowledgements

We thank the research coordinators who participated at various stages of data collection for this study, including Shilpa Karanjit, Ariel Watts, Holly Cirri, and Ann Onyenwoke. We also thank the NC Translational and Clinical Sciences Institute (NCTraCs) for funding via Pilot Grant UNCSUR11609. M.D. was supported by NIH training grant T32 DK007634. E.S.B and G.Z. were funded by grants from the National Institute of Environmental Health Sciences (P42 ES027704), the National Institute of General Medical Sciences (R01 GM141277 and RM1 GM145416), and a cooperative agreement with the Environmental Protection Agency (STAR RD 84003201). C.M.T. is funded by the National Institute of General Medical Sciences of the National Institutes of Health under award number R35GM119438 and R35GM149222. We also acknowledge the computing resources provided by North Carolina State University High Performance Computing Services Core Facility (RRID:SCR_022168) and Samantha Kisthardt for the assistance in annotation of Stickland genes.

## Author Contributions

Conceptualization, M.K.D, S.K.M, A.S.G, and C.M.T; Data curation, A.S.M and G.Z; Formal Analysis, A.S.M and G.Z ; Funding acquisition, M.K.D, S.K.M, A.S.G, and C.M.T; Investigation, A.S.M, G.Z, M.K.D, S.K.M, A.S.G, E.S.B, and C.M.T; Methodology, A.S.M, G.Z, M.K.D, S.K.M, A.S.G, E.S.B, and C.M.T; Project administration, M.K.D, S.K.M, A.S.G, E.S.B, and C.M.T; Resources, M.K.D, S.K.M, A.S.G, E.S.B, and C.M.T; Software, A.S.M, G.Z, and E.S.B ; Supervision, M.K.D, C.M.T and E.S.B; Validation, A.S.M, G.Z, E.S.B, and C.M.T; Visualization, ASM; Writing – original draft A.S.M, M.K.D, and C.M.T; Writing – review & editing, A.S.M, G.Z, M.K.D, S.K.M, A.S.G, E.S.B, and C.M.T

## Declaration of interests

C.M.T. consults for Vedanta Biosciences, Inc., Summit Therapeutics, and Ferring Pharmaceuticals, Inc. and is on the Scientific Advisory Board for Ancilia Biosciences.

## Declaration of generative AI and AI-assisted technologies in the writing process

During the preparation of this work the author(s) used Paperpal for assistance with grammar. After using this tool, the author(s) reviewed and edited the content as needed and take(s) full responsibility for the content of the publication.

## STAR Methods

### Patient enrollment

We enrolled all consenting patients undergoing FMT for multiply rCDI at the University of North Carolina from January to December 2017 in a prospective registry. Multiply rCDI was defined as at least the third episode of *C. difficile* infection. There were no exclusion criteria for participation in the registry specifically, though subjects were by definition undergoing FMT under the care of a physician who judged the benefits to outweigh the risks. The registry collected data on clinical characteristics and outcomes, and patient stool samples from pre-FMT (n=16), and 2 weeks (n=11), 2 months (n=10), and 6 months (n=8) post-FMT. Informed written consent was obtained from recipients. Stools samples were collected and de-identified by the research team. In raw data files individuals are given a subject identifier of R[1-15], and timepoints are labeled 1, 2, 3, 4 respective to sampling timepoints listed previously. Subjects’ clinical data was collected by research personnel via in-person or telephone interviews as well as medical record review and discussion with treating clinicians, and managed using REDCap electronic data capture tools hosted a UNC. Data collection timepoints were at enrollment (prior to FMT), and a post-FMT clinical questionnaire at two time points during the first 6 months post-FMT, and at 12 months post-FMT (Appendix A), to detect recurrence as well as adverse events. The study was approved by the UNC Institutional Review Board (#16-2283).

### Metagenomic analysis

Samples were processed and sequenced at CoreBiome (now Diversigen).

#### Library Preparation & Sequencing

Libraries were prepared with a procedure adapted from the Nextera Library Prep kit (Illumina). Libraries were sequenced using single-end 1 x 150 reads with a NextSeq 500/550 High Output v2 kit (Illumina).

#### DNA extraction

Samples were extracted with the MO Bio PowerFecal kit (Qiagen) automated for high throughput on QiaCube (Qiagen). The manufacturer’s instructions were followed with bead beating in 0.1mm glass bead plates.

#### DNA quantification

Samples were quantified with Qiant-iT Picogreen dsDNA Assay (Invitrogen). Shallow shotgun sequencing of fecal DNA on the Illumina Nextseq platform was performed by CoreBiome, now Diversigen. Raw reads were filtered for quality and reads that map to a human genome were removed using KneadData v0.12.0. The relative abundances of each bacterial phylum and species member of pre-FMT and post-FMT fecal samples were determined using Metaphlan version 4.0.6 ^62^ using the vOct22 CHOCOPhlanSGB database with default settings and annotated with Genome Taxonomy Database (GTDB) notation. Microbial genes were determined using Humann version 3.7 ^63^ using the vOct22 CHOCOPhlanSGB database. To account for decreased depth due to shallow shotgun sequencing the threshold for a nucleotide match required to call a gene present was reduced from 50% coverage to 20% coverage. UniRef90 was used as the reference database for the translated alignment search.

#### Mapped read analysis

Alpha and beta diversity were examined using the R package vegan v2.6- 4 ^64^. BSH genes were selected by UniRef90 Protein ID queries for “bile salt hydrolase” or “chololylglycine hydrolase” with further manual curation (Table S|raw data|). MUSCLE alignment of BSH genes for loop structure identification was performed using the R package muscle v 3.40.0 ^65^. Glycine-preferring genes encode the GXG motif at the specificity loop while Taurine-preferring encode an SRX motif ^46^. For species level analysis of BSH genes each species is represented by the sum total counts per million (CPM) of every BSH gene encoded by that species. Other bile acid altering enzymes were selected by UniRef90 Protein ID queries for “bile”, “hydroxysteroid”, “hydroxycholanate”, “choloylglycine”, and “bai” with further manual curation (Table S|raw data|). Antimicrobial resistance genes were selected by UniRef90 Protein ID queries for both “resistance” and the name of an antibiotic with further manual curation (Table S|raw data|). Microbial pathway analyses were performed using UniPathway grouping of UniRef90 genes identified by Humann. Amino acid metabolism pathways were aggregated as the sum CPM of all pathways that synthesize or degrade each amino acid. Species were clustered using complete linkage clustering with Euclidian distances by the R package pheatmap v1.0.1266.

### Untargeted metabolomic analyses

Metabolomic profiling analysis was performed by Metabolon (Durham, NC) in the following way.

#### Sample accessioning

Following receipt, samples were inventoried and immediately stored at - 80°C. Each sample received was accessioned into the Metabolon LIMS system and was assigned by the LIMS a unique identifier that was associated with the original source identifier only. This identifier was used to track all sample handling, tasks, results, etc. The samples (and all derived aliquots) were tracked by the LIMS system. All portions of any sample were automatically assigned their own unique identifiers by the LIMS when a new task was created; the relationship of these samples was also tracked. All samples were maintained at -80°C until processed.

#### Sample Preparation

Samples were prepared using the automated MicroLab STAR® system from Hamilton Company. Several recovery standards were added prior to the first step in the extraction process for QC purposes. To remove protein, dissociate small molecules bound to protein or trapped in the precipitated protein matrix, and to recover chemically diverse metabolites, proteins were precipitated with methanol under vigorous shaking for 2 min (Glen Mills GenoGrinder 2000) followed by centrifugation. The resulting extract was divided into 5 fractions: 2 for analysis by 2 separate reverse phase (RP)/UPLC-MS/MS methods with positive ion mode electrospray ionization (ESI), one for analysis by RP/UPLC-MS/MS with negative ion mode ESI, one for analysis by HILIC/UPLC-MS/MS with negative ion mode ESI, and one sample was reserved for backup. Samples were placed briefly on a TurboVap® (Zymark) to remove the organic solvent. The sample extracts were stored overnight under nitrogen before preparation for analysis.

#### QA/QC

Several types of controls were analyzed in concert with the experimental samples: a pooled matrix sample generated by taking a small volume of each experimental sample (or alternatively, use of a pool of well-characterized human plasma) served as a technical replicate throughout the data set; extracted water samples served as process blanks; and a cocktail of QC standards that were carefully chosen not to interfere with the measurement of endogenous compounds were spiked into every analyzed sample, allowed instrument performance monitoring and aided chromatographic alignment. Instrument variability was determined by calculating the median relative standard deviation (RSD) for the standards that were added to each sample prior to injection into the mass spectrometers. Overall process variability was determined by calculating the median RSD for all endogenous metabolites (i.e., non-instrument standards) present in 100% of the pooled matrix samples. Experimental samples were randomized across the platform run with QC samples spaced evenly among the injections.

#### Ultrahigh Performance Liquid Chromatography-Tandem Mass Spectroscopy (UPLC-MS/MS)

All methods utilized a Waters ACQUITY ultra-performance liquid chromatography (UPLC) and a Thermo Scientific Q-Exactive high resolution/accurate mass spectrometer interfaced with a heated electrospray ionization (HESI-II) source and Orbitrap mass analyzer operated at 35,000 mass resolution. The sample extract was dried then reconstituted in solvents compatible to each of the four methods. Each reconstitution solvent contained a series of standards at fixed concentrations to ensure injection and chromatographic consistency. One aliquot was analyzed using acidic positive ion conditions, chromatographically optimized for more hydrophilic compounds. In this method, the extract was gradient eluted from a C18 column (Waters UPLC BEH C18-2.1x100 mm, 1.7 µm) using water and methanol, containing 0.05% perfluoropentanoic acid (PFPA) and 0.1% formic acid (FA). Another aliquot was also analyzed using acidic positive ion conditions, however it was chromatographically optimized for more hydrophobic compounds. In this method, the extract was gradient eluted from the same afore mentioned C18 column using methanol, acetonitrile, water, 0.05% PFPA and 0.01% FA and was operated at an overall higher organic content. Another aliquot was analyzed using basic negative ion optimized conditions using a separate dedicated C18 column. The basic extracts were gradient eluted from the column using methanol and water, however with 6.5mM Ammonium Bicarbonate at pH 8. The fourth aliquot was analyzed via negative ionization following elution from a HILIC column (Waters UPLC BEH Amide 2.1x150 mm, 1.7 µm) using a gradient consisting of water and acetonitrile with 10mM Ammonium Formate, pH 10.8. The MS analysis alternated between MS and data-dependent MS^n^ scans using dynamic exclusion. The scan range varied slighted between methods but covered 70-1000 m/z. Raw data files are archived and extracted as described below.

#### Data Extraction and Compound Identification

Raw data was extracted, peak-identified and QC processed using Metabolon’s hardware and software. These systems are built on a web-service platform utilizing Microsoft’s .NET technologies, which run on high-performance application servers and fiber-channel storage arrays in clusters to provide active failover and load-balancing. Compounds were identified by comparison to library entries of purified standards or recurrent unknown entities. Metabolon maintains a library based on authenticated standards that contains the retention time/index (RI), mass to charge ratio (*m/z)*, and chromatographic data (including MS/MS spectral data) on all molecules present in the library. Furthermore, biochemical identifications are based on three criteria: 1. retention index within a narrow RI window of the proposed identification, 2. accurate mass match to the library +/- 10 ppm, and 3. the MS/MS forward and reverse scores between the experimental data and authentic standards. The MS/MS scores are based on a comparison of the ions present in the experimental spectrum to the ions present in the library spectrum. While there may be similarities between these molecules based on one of these factors, the use of all three data points can be utilized to distinguish and differentiate biochemicals. More than 3300 commercially available purified standard compounds have been acquired and registered into LIMS for analysis on all platforms for determination of their analytical characteristics. Additional mass spectral entries have been created for structurally unnamed biochemicals, which have been identified by virtue of their recurrent nature (both chromatographic and mass spectral). These compounds have the potential to be identified by future acquisition of a matching purified standard or by classical structural analysis.

#### Curation

A variety of curation procedures were carried out to ensure that a high quality data set was made available for statistical analysis and data interpretation. The QC and curation processes were designed to ensure accurate and consistent identification of true chemical entities, and to remove those representing system artifacts, mis-assignments, and background noise. Metabolon data analysts use proprietary visualization and interpretation software to confirm the consistency of peak identification among the various samples. Library matches for each compound were checked for each sample and corrected if necessary.

#### Metabolite Quantification and Data Normalization

Peaks were quantified using area-under-the-curve. For studies spanning multiple days, a data normalization step was performed to correct variation resulting from instrument inter-day tuning differences. Essentially, each compound was corrected in run-day blocks by registering the medians to equal one (1.00) and normalizing each data point proportionately (termed the “block correction”). For studies that did not require more than one day of analysis, no normalization is necessary, other than for purposes of data visualization. In certain instances, biochemical data may have been normalized to an additional factor (e.g., cell counts, total protein as determined by Bradford assay, osmolality, etc.) to account for differences in metabolite levels due to differences in the amount of material present in each sample.

### Targeted metabolomic analysis

#### BA and MCBA Extraction

To extract the bile acids (BAs) from the 45 fecal samples, approximately 100 mg of each fecal sample was aliquoted into a 2 mL Omni International microtubes containing 2.38 mm metal beads (Kennesaw, Georgia, catalog number 19-620). The slight variations in fecal mass were normalized to a 1 mg:8 μL ratio by adding eight times the mass (mg) of a pre-extraction buffer (μL) consisting of 1:1 methanol (MeOH):acetonitrile (ACN) with 3 mM phosphate. For example, for 100.1 mg of fecal sample, 800.8 μL of extraction buffer was added. For the extraction buffer, Optima™ LC/MS grade methanol (catalog number 67-56- 1) and acetonitrile (catalog number 75-05-8) were purchased from Fisher Chemical (Pittsburgh, PA) and the phosphate was potassium phosphate monobasic from Supelco (Bellefonte, PA, catalog number 57618).

A C13 heavy labeled BA internal standard mix was also prepared for the study by combining two BA mixes purchased from Cambridge Isotope Laboratories (CIL, Tewksbury, MA), a labeled conjugated BA mix (MSK-BA1-US-1) and a labeled unconjugated BA mix (MSK-BA1-US-2). Each mix vial was resuspended in 250 μL of a 1:1 mixture of MeOH:H_2_O (catalog numbers 67-56-1 and 7732-18-5). The contents of both vials were then combined, resulting in a 500 μL BA internal standard mix with a concentration of ∼200 μM for each BA (0.08 mg/mL MSK-BA1-US-1 and 0.1 mg/mL MSK-BA2-US-2). Subsequently, 5 μL of the internal BA standard was spiked into all 2 mL Omni International microtubes including calibrations (quality controls) and blanks prior to all extraction steps. Homogenization of the fecal sample/pre-extraction buffer mixtures was achieved using a Fisherbrand™ Bead Mill 24 Homogenizer (Hampton, NH, catalog number 15-340-163), followed by centrifugation at 13,000LJ×LJg at 4 °C for 10 min in an Eppendorf 5810R centrifuge (Hamburg, Germany, catalog number 022625501). Following centrifugation, a 500 μL aliquot of the supernatant was taken from the tube and stored in a 1.5 mL Thermo Scientific microcentrifuge tube (San Diego, CA, catalog number 3451) for the subsequent extraction steps. Only a 150 uL portion of the pre-extracted supernatant was then aliquoted into a different 1.5 mL Thermo Scientific microcentrifuge tube (San Diego, CA, catalog number 3451) and combined at a ratio of 1:1 volume/volume (v/v) with 150 μL of Optima™ LC/MS grade MeOH (Fisher Chemical, Pittsburgh, PA, catalog number 67-56-1) and shaken on an Eppendorf MixMate ^®^ (Hamburg, Germany, 5353000529) at 600 rpm and room temperature for 20 mins. The resulting mixtures were then filtered into with a Millipore Ultrafree Centrifugal PTFE filter with its own tube (Jaffrey, NH, catalog number UFC30LG25) and centrifuged at 10,000LJ×LJg at 4 °C for 1LJmin. The filtrate was diluted 2:1 volume/volume (v/v) with H_2_O (Fisher Chemical, Pittsburgh, PA, catalog number 7732-18-5) in Agilent high recovery vials (Santa Clara, CA, catalog number 5188-6591) and then injected into the LC column for LC-IMS-MS analyses.

Blanks and calibration samples were also utilized for the study, with the calibration samples also serving to assess extraction efficiency. An unlabeled BA standard mix was prepared for the calibration samples. This unlabeled BA mix consisted of 16 unlabeled BAs purchased from Cambridge Isotope Laboratories (CIL, Tewksbury, MA) including conjugated BAs in vial 1 (catalog number MSK-BA1) and unconjugated BAs in vial 2 (catalog number MSK-BA2). To prepare the standard BA mix, each vial was resuspended in 500 μL of a 1:1 mixture of MeOH:H_2_O (catalog numbers 67-56-1 and 7732-18-5), and the contents of both vials were combined, resulting in a total of 1 mL with a concentration of approximately 100 μM for each BA (0.04 mg/mL MSK-BA1 and 0.05 mg/mL MSK-BA2). The standard mix was further diluted into varies concentrations ranging from 3.2 nM to 50 μM (3.2 nM, 16 nM, 80 nM, 400 nM, 2 μM, 10 μM, and 50 μM). For the calibration curves, 100 μL of standard mix at each concentration was utilized. Additionally, both an extraction blank, containing 100 µL of the 1:1 MeOH:H_2_O resuspension solvent for the BA standard mix and a method blank with all solvents but no fecal material were created. To replicate the sample preparation steps as much as possible, both the calibration samples and blanks were diluted to a 1:8 volume/volume (v/v) with the pre-extraction buffer then taken through all the extraction steps.

#### LC-IMS-MS BA and MCBA Lipidomic Analyses

Simultaneous liquid chromatography, ion mobility spectrometry and mass spectrometry (LC-IMS-MS) analyses were performed utilizing an Agilent 1290 Infinity UPLC (Santa Clara, CA) system coupled with an Agilent 6560 IM-QTOF MS instrument (Santa Clara, CA).^67,68^ Prior to sample analysis, 6 μL of the solvent blank was initially evaluated to ensure no contaminants were observed. Instrumental performance and evaluation of the separation efficiency were assessed using both the unlabeled BA standard mix created above from CIL samples and a BA standard mix purchased from Cayman Chemical (Ann Arbor, MI, catalog number 33505). Following the performance checks, the 45 final extracted fecal samples, along with the 7 calibration samples and 2 blanks samples, were randomized and assessed by LC-IMS-CID-MS. Solvent blanks were also conducted every 9 sample injections to evaluate carry-over. At the end of the samples runs, the BA standard mixes were rerun to check instrument performance.

The LC separation was slightly modified from our previously described methods to achieve higher resolution of the conjugated BA.^45,46^ In this study, the Restek Raptor C18 column (1.7 μm, 2.1 × 50 mm) (Bellefonte, PA, catalog number 9304A52) was again used with a column temperature of 60LJ°C.^45,46^ The column flow rate was set to 0.5 mL/min and the LC runtime was 13.5 min (11 min of acquired gradient time and 2.5 min for column wash and re-equilibration). Specific details of the LC gradient are provided in Table S7, where mobile phase A consisted of 5 mM ammonium acetate in H_2_O (catalog number 7732-18-5), and mobile phase B was a 1:1 mixture of MeOH (catalog number 67-56-1) and ACN (catalog number 75-05-8).

All mobile phase solvents were LC/MS grade and purchased from Fisher Chemical (Pittsburgh, PA) including the ammonium acetate (catalog number 631-61-8).

The LC-IMS-CID-MS platform had been characterized in previous studies.^67–69^ Mass calibration was performed prior to sample analysis using electrospray ionization (ESI) in negative ion mode and Agilent ESI tune mix (Santa Clara, CA, catalog number G1969-85000). The instrument source parameters were set as follows: gas temperature at 325 °C, drying gas at 10 L/min, nebulizer at 40 psi, sheath gas temperature at 275 °C, and sheath gas flow at 12 L/min. Following ionization, the ions traveled through a single-bore glass capillary, were focused into a high-pressure funnel, and then accumulated in a trap funnel. Ions were then pulsed into an approximately 78 cm long IMS drift tube with 3.95 Torr of nitrogen drift gas, using Hadamard transform multiplexing. Here, a 4-bit pseudo-random pulsing sequence was utilized for packet ejection out of the trapping funnel and a trap fill time of 3.9 ms and release time of 100 μs.^70^ After exiting the drift tube, ions were refocused in a rear ion funnel and analyzed in a quadrupole time-of-flight mass spectrometer. Data were collected for a mass range of 50 to 1700LJ*m/z*. All LC retention time, IMS drift time, and *m/z* multiplexed data was collated into a raw .d file.

#### Skyline Statistical Analyses

The LC-IMS-MS .d files were demultiplexed with the PNNL- Preprocessor 4.0 at a moving average of 3, minimum pulse coverage of 100%, and signal intensity threshold of 20 counts, resulting in (.DeMP.d) files.^71^ The demultiplexed data files were single-field–calibrated for drift time to CCS conversions using Agilent ESI tuning mix data in the Agilent MassHunter IM-MS Browser 10.0 software.^67^ These files were subsequently uploaded to the open-source software Skyline 23.1^72^ (MacCoss lab, Seattle, Washington, release date 9/24/23) for peak picking. A BA/MCBA library with 208 entries was utilized for the BA and MCBA identifications. Specifically, the library included 35 unique BA standards purchased from Cambridge Isotope Laboratories (CIL, Tewksbury, MA) and Cayman Chemical (Ann Arbor, MI), and an additional 173 MCBAs.^45^ The synthesis and evaluation of MCBA standards were conducted by Quinn *et al*.^61^, employing the methodologies established by Ezawa *et al*.^73^ The BAs and MCBAs as listed in Table S8 and the full library can be found at https://panoramaweb.org/rT2GQI.url in Panorama Public ^74,75^ (MacCoss lab, Seattle, WA). Following identification in Skyline, each BA, MCBA and their corresponding peak area under the curve (AUC) in each sample was exported with mass errors to Microsoft Excel. Mass errors greater than 10 ppm were removed from further data processing.

### Random Forest Analysis

Important metabolites were defined as having an MDA>0 as determined using random forest analysis via the R package randomForest v4.7-1.1 ^76^. For all random forest analyses the number of trees was set to 1000. The number of variables randomly sampled as candidates at each split was left at the default setting of the square root of the total number of variables.

### Microbiome and metabolome correlation analysis

Repeated measures correlation (rmcorr) between microbiome and metabolome were performed using the R package rmcorr v0.6.0 ^47^. Correlations were done between taxa and biochemicals that were significantly different between timepoints or after FMT by linear mixed models with Bonferroni correction (q<0.05) unless otherwise noted. Correlations deemed significant (p<0.05) after Benjamini and Hochberg correction are marked with an asterisks after complete linkage clustering using the Pearson correlation coefficients of the repeated measure correlations with the R package pheatmap v1.0.12 where applicable ^66^.

### Statistical analysis

Significant correlations were determined by linear model with Benjamini and Hochberg correction on the correlation values using the R package rmcorr v0.6.0 and stats v4.2.2 ^47^. Significant differences between timepoints stratified by participant in NMDS were determined by Adonis using the R package pairwiseAdonis v0.4.1 ^77^. Linear mixed models were generated using MaAsLin2 v1.15.1 ^78^ with separate models to determine significance between pre-FMT and all post-FMT timepoints as well as between pre-FMT and 2 week, 2 month, and 6 month timepoints individually. Data was log transformed prior to linear modelling. To account for longitudinal sampling of the same individual, the subject ID for each sample was declared as a random effect. Linear models were generated for family level taxonomic data, microbial genes, microbial pathway abundances, untargeted metabolomics, and targeted metabolomics. Wilcoxon signed rank tests with Holm correction to determine differences in CPM of genes & pathways between timepoints was performed using the R packages rstatix v0.7.2 and ggpubr v0.6.0 ^79^. Boxplots depict inter-quartile range (IQR) with whiskers depicting the most extreme value or 1.5X IQR, whichever is lower. For statistical tests requiring paired samples involving the failed FMT, R9, the timepoint before any FMT, R9_1, was used as the baseline for comparison. Plotting and data organization was performed using tidyverse ^80^. All other operations were performed in base R v4.2.2 ^81^.

### Data and code availability

Raw sequences from shallow shotgun sequencing have been deposited in the Sequence Read Archive (SRA) under BioProject ID PRJNA1055134. Data acquired from targeted metabolomics have been deposited in MASSive under ftp://MSV000093844@massive.ucsd.edu Source data and code required for statistical analysis and figure generation have been deposited in GitHub and are available at https://github.com/asmcmill/FMT-Manuscript, doi:___. Source data and statistical analyses are also provided with Mendeley data: Reserved DOI: 10.17632/8s9mdf8xrj.1. Other data and biological materials are available from the corresponding author upon reasonable requests.

## Notes

https://github.com/asmcmill/FMT-Manuscript

