## Supplementary figures and images for "Metagenomic, metabolomic, and lipidomic shifts associated with fecal microbiota transplantation for recurrent *Clostridioides difficile* infection"

### Figure S1

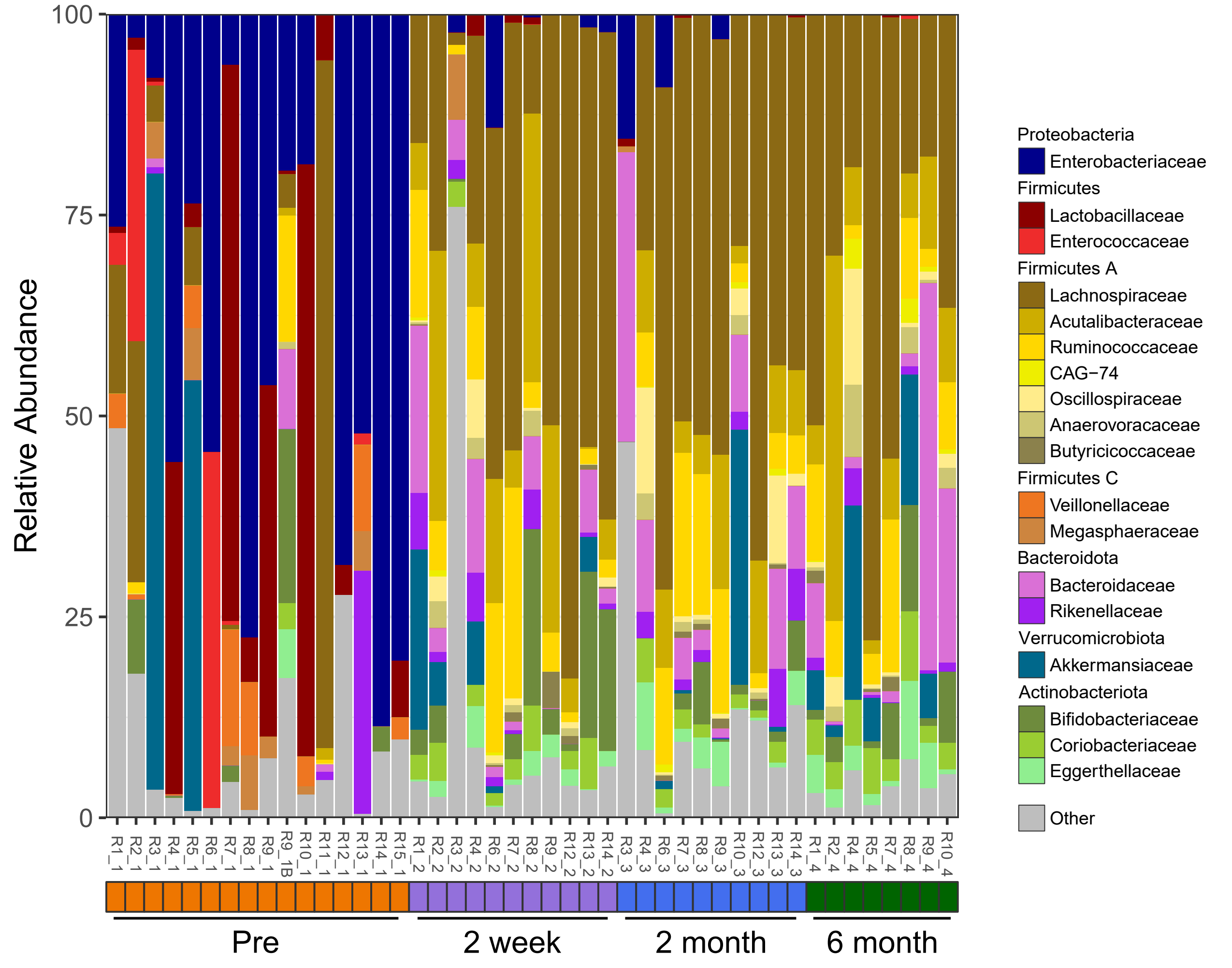

### Figure S2

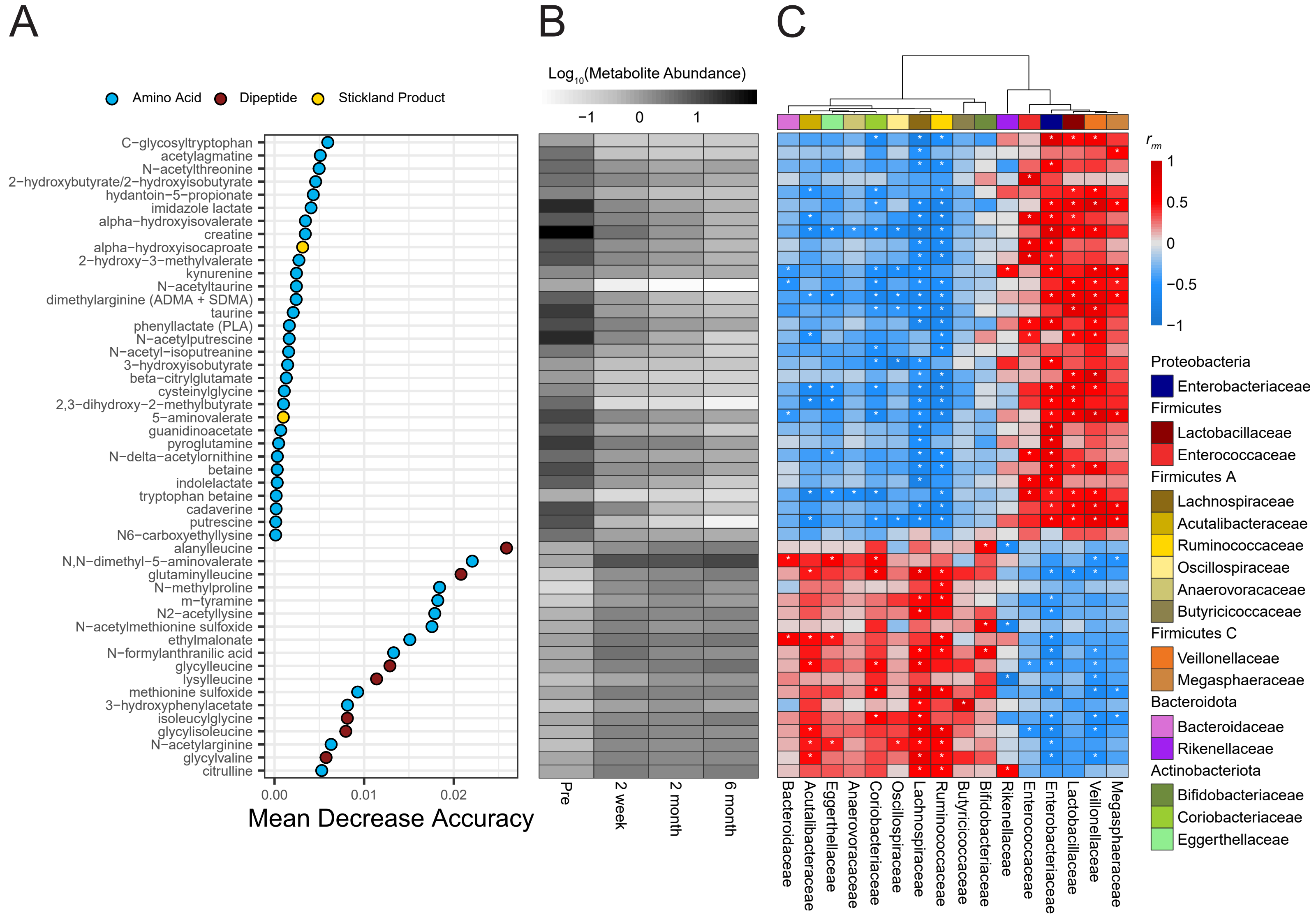

### Figure S3

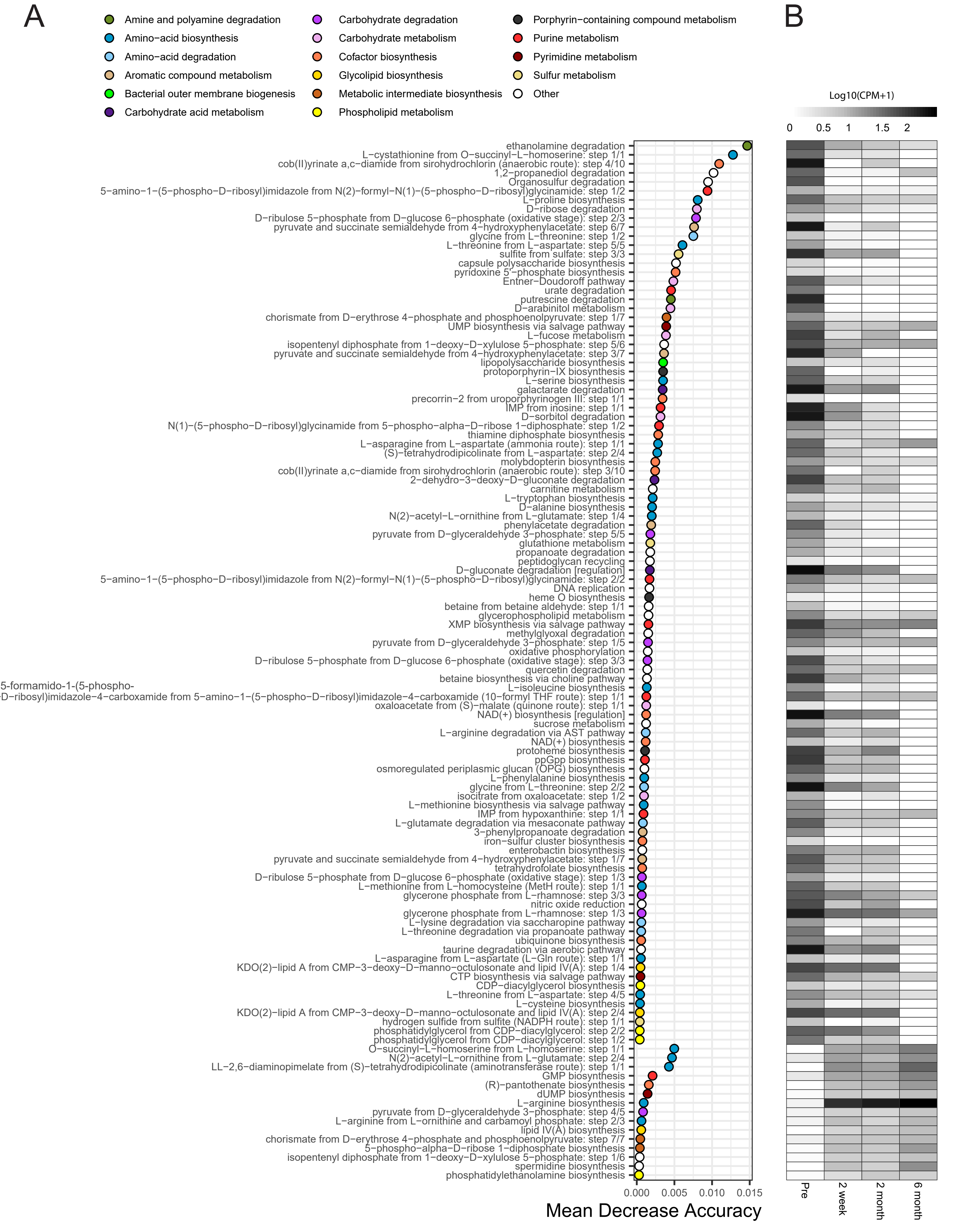

### Figure S4

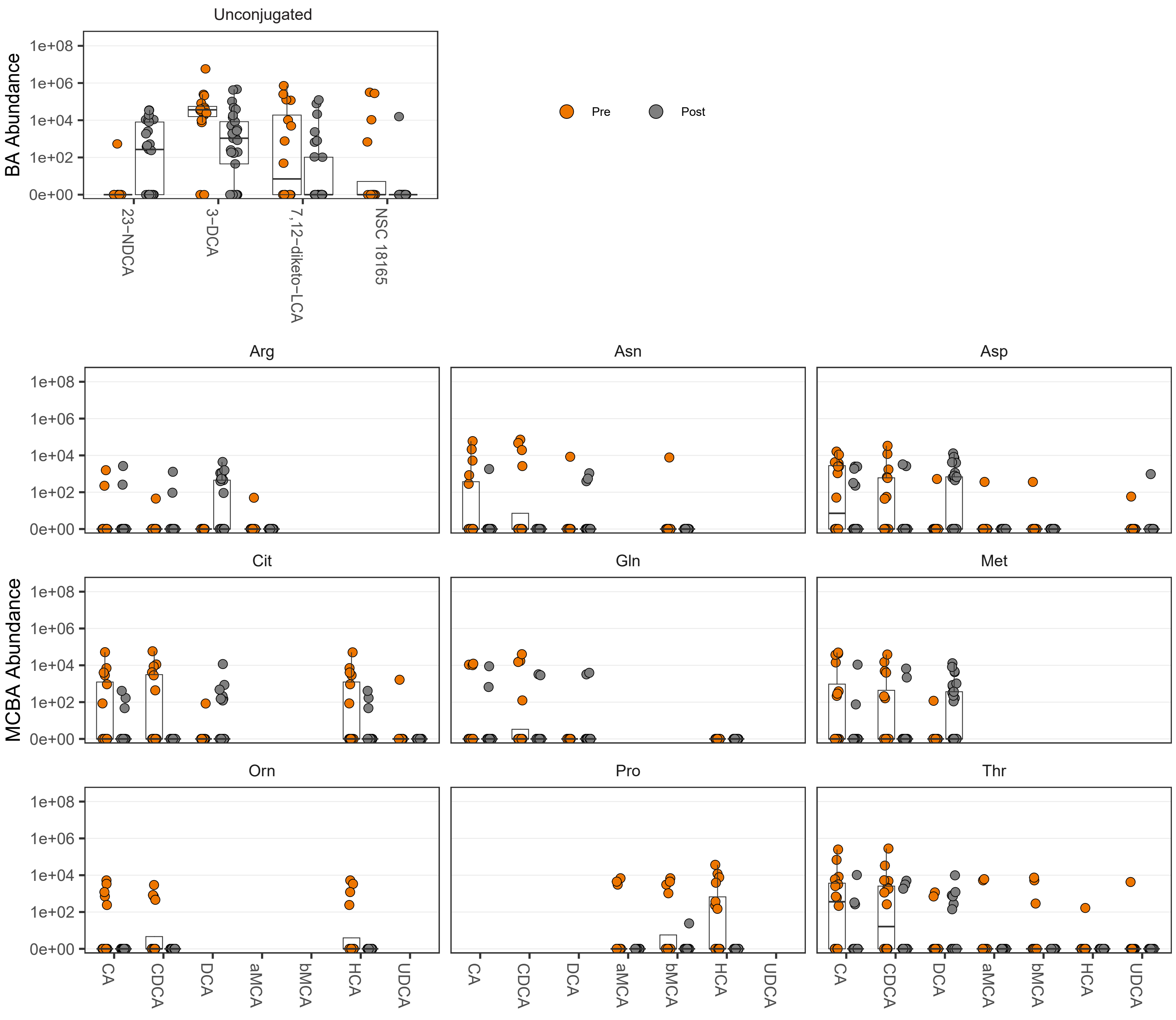

### Figure S5

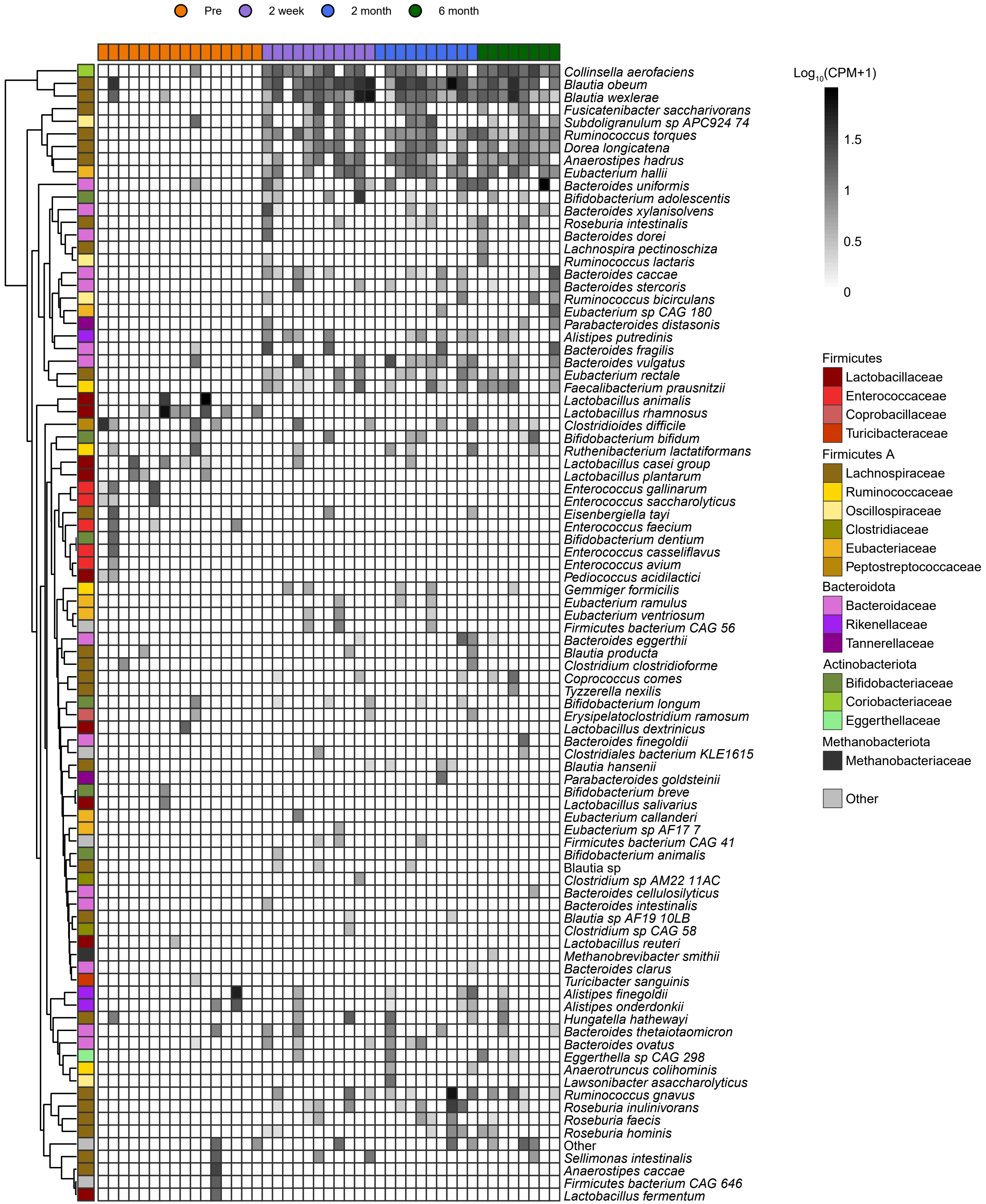

### Figure S6

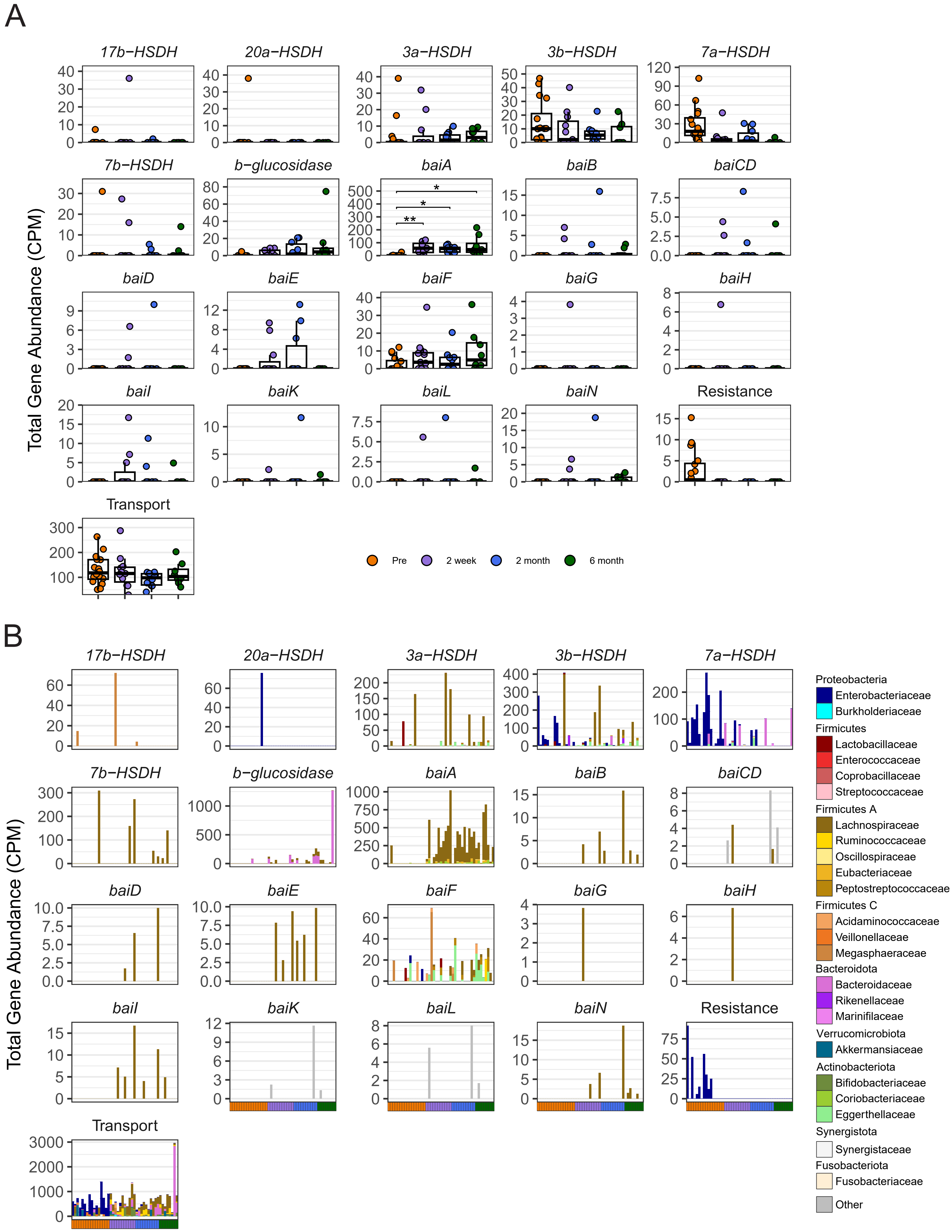
